# Drought-induced changes in sex-specific floral chemistry reshape interactions with a nursery pollinator across life stages

**DOI:** 10.64898/2026.10.22.760926

**Authors:** Karin Schrieber, Reza Esmaeilnia Talemi, Kaya Zill, Henry Berndt, Alexander Weinhold, Alexandra Erfmeier, Thomas Stegemann

## Abstract

Nursery pollination systems provide unique models for studying the chemical ecology of plant–insect interactions, as specialized insects act as both pollinators and seed predators across life stages. How climatic stress alters the plant chemistry underlying these interactions remains unknown.

To address this gap, we use the dioecious plant species *Silene latifolia* and its nursery pollinator *Hadena bicruris*. We investigated how water limitation affects visual floral traits, scent and nectar chemistry in male and female plants along with associated moth behaviour, as well as female seed chemistry along with associated larval performance.

Water limitation broadly compromised floral traits, with particularly pronounced reductions in nectar metabolites in male plants eroding sexual dimorphism in nectar chemistry. Correspondingly, moths preferred well-watered flowers for oviposition and lost their adaptive preference for female flowers under water limitation, while feeding behaviour remained unaffected. Water limitation also compromised seed traits, including reductions in defence-associated metabolites. In parallel, larvae grew faster on seeds from water-limited plants despite consuming similar amounts.

Our findings provide first evidence that climatic stress broadly compromises the multimodal phenotypes underlying nursery pollination, particularly chemical traits, and that these changes coincide with contrasting insect responses across life stages, with potential consequences for fitness and coevolution.

## 1 Introduction

Understanding how climate change reshapes the eco-evolutionary dynamics of mutualistic and antagonistic plant–insect interactions is a pressing challenge across biological disciplines. It requires mechanistic insight into how stress-induced changes in plant traits, particularly their chemical components, affect communication, resource use and ultimately fitness in both partners (Cantwell-Jones *et al*. 2024; Peralta *et al*. 2024). Nursery pollination systems offer unique, yet untapped opportunities for integrative assessments of these processes, as the same specialized insect pollinates its host plant as an adult and consumes its reproductive tissues (seeds, ovaries, anthers) as a larva.

Climatic stress can disrupt plant mutualistic interactions with pollinators by altering the complex floral signals underlying communication and rewards sustaining pollinators: Plants advertise their flowers through spatial visual signals such as shape, size, and arrangement (Dafni *et al*. 1997), colours determined by pigment composition and petal surface structures (van der Kooi *et al*. 2021), and scent profiles defined by the composition of numerous volatile organic compounds (VOCs) (Mostafa *et al*. 2022). Beyond these signals, nectar provides pollinators with a readily accessible source of energy and nutrients while also containing bioactive compounds that promote pollinator health and modulate behaviour (Barberis *et al*. 2023; Nepi *et al*. 2018). All these traits can change quantitatively and qualitatively under climatic stress, with consequences for pollinator visitation and, ultimately, plant reproductive success (Alquichire-Rojas *et al*. 2024; Brunet *et al*. 2024). Yet, the multidimensional floral phenotype is rarely considered as a whole, despite potentially contrasting responses of its component traits to climatic stress, and studies linking these changes to pollinator behavioural decisions remain scarce (Brunet *et al*. 2024; Kuppler & Kotowska 2021).

In nursery pollination systems, interactions with specialized insect species depend not only on floral signals and rewards influencing adult behaviour and performance, but also on the nutritional quality and chemical defence of reproductive tissues consumed by their larvae (Dufay 2010). These interactions may impose particularly complex selective pressures on plant chemistry and potentially favour delicate chemical profiles that balance adult attraction and reward with defence against larval consumption. Despite this functional coupling, our general understanding of the chemical ecology of nursery pollination remains fragmentary, with research focusing predominantly on floral volatile signals mediating adult attraction (Cao *et al*. 2026; reviewed in Hossaert-McKey *et al*. 2010), while the chemistry of rewards for adults (i.e., nectar, pollen) and larval resources has received very little attention (see Gallet *et al*. 2007; Ibanez *et al*. 2009; Zill *et al*. 2025). Even less is known about how climatic stress reshapes the complex plant traits mediating these interconnected stages and how such changes affect insects across life stages. Studies of pollinating herbivores (adults pollinate, larvae feed on vegetative tissues) have provided compelling evidence that temperature-induced changes in plant traits can affect insect behaviour, herbivory and reciprocal selection (Rusman *et al*. 2025; Traine *et al*. 2024). Nursery pollination systems offer an even more tightly coupled context for investigating these processes, as both the mutualistic benefits and antagonistic costs directly affect plant reproduction. The few existing studies have focused primarily on direct temperature effects on nursery pollinator survival, development and reproductive opportunities, demonstrating that warming can alter the balance between plant and pollinator reproduction (van Kolfschoten *et al*. 2022; Krishnan *et al*. 2014; Wang *et al*. 2009). In contrast, how climatic stress affects nursery pollination through changes in host plant traits remains largely unexplored.

An additional dimension of these plant-mediated effects arises from dioecy, the separation of female and male reproductive functions among individuals, which is strikingly overrepresented among nursery pollination host plants (Dufaÿ & Anstett 2003; Labouche & Bernasconi 2010). Here, reproductive tissue consumption by larvae is limited to individuals of one sex, while pollination by adults is maintained across the entire population. This asymmetric distribution of interaction costs and benefits between the plant sexes generates sex-specific selection on traits mediating pollinator attraction, reward and feeding resistance promoting sexual dimorphism (Dufaÿ & Anstett 2003; Labouche & Bernasconi 2010). Male and female plants of diecious species can also differ in their physiological responses to climatic stress (Chen *et al*. 2026), potentially altering the magnitude or direction of sexual dimorphism in floral traits. However, whether climatic stress reshapes these functionally important differences, and with what consequences for interactions with nursery pollinators, remains an open question.

A suitable system to approach these open questions consists of the dioecious plant species *Silene latifolia* POIR (Caryophyllaceae), a well-established model in ecology and evolutionary biology (Karrenberg *et al*. 2026), and its nursery moth pollinator *Hadena bicruris* HUFNAGEL (Lepidoptera: Noctuidae). In this system, *H. bicruris* larvae consume developing seeds, restricting antagonistic costs to female plants. Previous research has characterized pronounced sexual dimorphism in *S. latifolia* (e.g., Schrieber *et al*. 2021; Waelti *et al*. 2009) and investigated various aspects of the ecology of its interaction with *H. bicruris* (Karrenberg *et al*. 2026; Prieto-Benítez *et al*. 2017). However, the functional traits mediating this interaction have been studied predominantly in the context of visual traits and floral scent communication (e.g., Dötterl *et al*. 2006; Dötterl & Jürgens 2005; Waelti *et al*. 2009). In our recent work, we extended this perspective to nectar chemistry (Zill *et al*. 2025), whereas the chemistry of seeds as the larval resource remains virtually unexplored. Here, we aim to resolve how integrated *S. latifolia* phenotypes respond to climatic stress in a sex-specific manner and affect interactions with *H. bicruris* across successive life stages.

To this end, we exposed female and male plants from five central European populations to controlled water limitation, assessing visual floral traits, floral scent, nectar chemistry and moth behaviour alongside with seed production, seed chemistry, larval consumption and performance. This integrative approach allowed us to address three questions: 1) Does water limitation reshape multimodal floral signalling and associated interactions with moths? (2) Do male and female plants differ in their responses to water limitation, thereby altering sexual dimorphism and sex-specific interactions with moths? (3) Does water limitation reshape seed provisioning and defence, with consequences for larval consumption and performance?

## 2 Material & Methods

### 2.1 Plant and pollinator study species

*Silene latifolia* (White Campion, ***Figure 1a***) is a perennial herb typical of ruderal habitats. Its flowers exhibit a characteristic moth-pollination syndrome, with large, funnel-shaped petals (Dafni et al., 1997) that open at dusk. Bright white petal colours arise from the intense, non-selective backscattering of visible light by specialized petal surface structures and epidermal cell architecture (van der Kooi *et al*. 2016). Upon opening, flowers emit a scent bouquet of about 70 VOCs, which is dominated by lilac aldehydes (Dötterl & Jürgens 2005). Scent emission peaks during early night and coincides with the presentation of large nectar rewards, which comprise numerous nutritious and potentially behaviourally active primary and secondary metabolites (Zill *et al*. 2025). *S. latifolia* exhibits a dioecious reproductive system, with pronounced sexual dimorphisms. In line with the extension of Bateman’s principle to plant ecology, many studies reported that male *S. latifolia* flowers are generally more attractive to pollinators (e.g., Waelti *et al*. 2009). However, variation in the magnitude and even the direction of these sexual differences among studies, and among populations within studies, challenges the assumption of such fixed evolutionary trajectories and instead points to locally driven, context-dependent trait evolution (Dötterl & Jürgens 2005; Schrieber *et al*. 2019, 2021; Yu *et al*. 2011; Zill *et al*. 2025). In Europe, *S. latifolia* flowers are visited by diurnal generalist pollinators and crepuscular – nocturnal insects.

**Figure 1:**
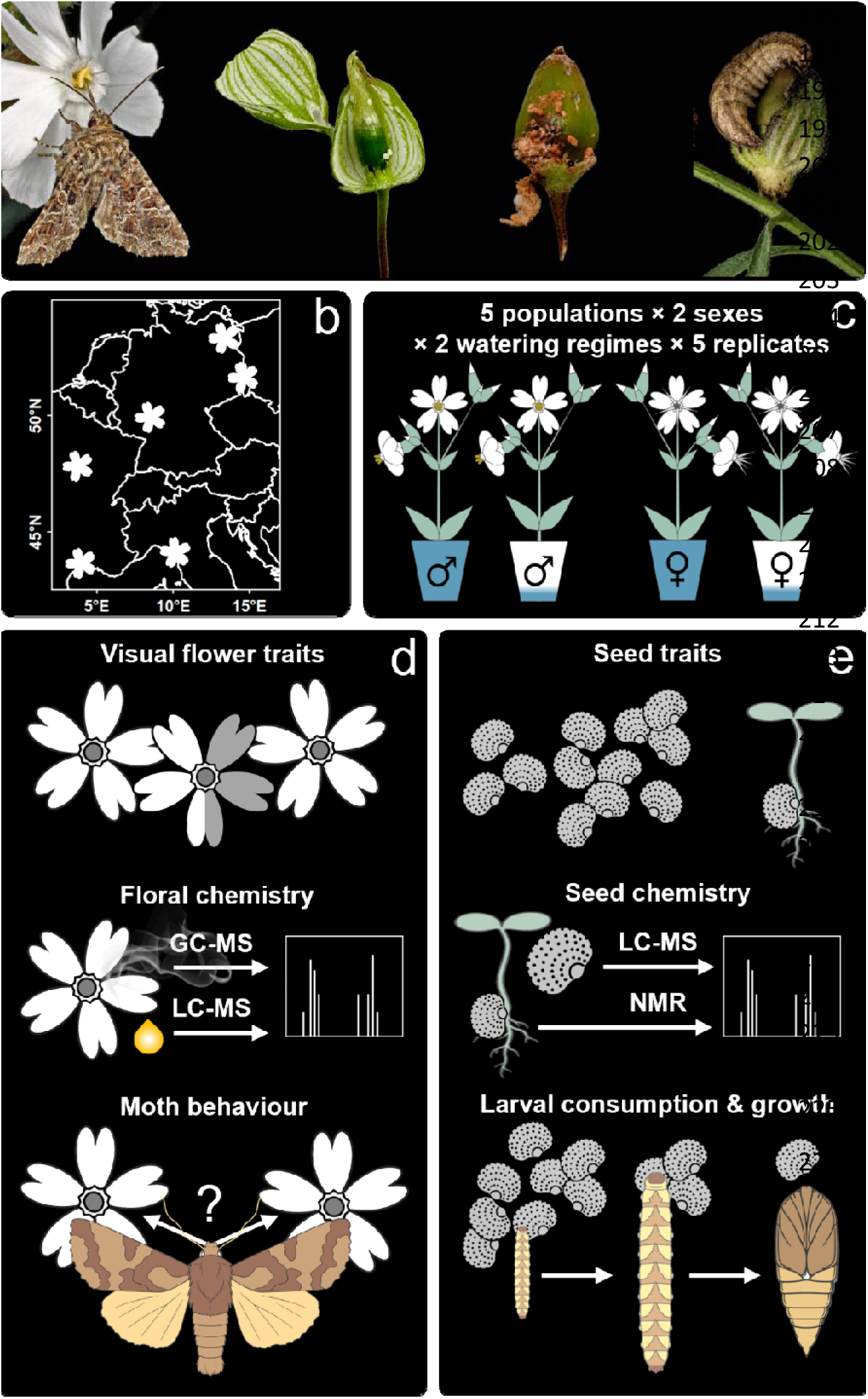
Overview of the experimental-setup and acquired 199-traits. (a) Plant (Silene 200-latifolia) and insect (Hadena bicruris) 202-study species. Photos illustrate flower visitation by adult moths (© Zoran Bozovic, Saxifraga Foundation, CC BY NC-SA) and the development of their offspring from eggs to larvae feeding on seed capsules. (b) Geographic origin of the five plant populations and the H. bicruris laboratory population (moth symbol). (c) Experimental design. (d) Overview of traits acquired on flowers and moths. (e) Overview of traits acquired on seeds and larvae.

Among the latter *Hadena bicruris* (Campion owl, ***Figure 1a***) – a moth of the Noctuidae family is the most frequent visitor (Jürgens *et al*. 1996; Schrieber *et al*. 2021; Scopece *et al*. 2018; van Putten *et al*. 2007). Adult *H. bicruris* activity peaks at dusk between May and July, matching the flowering phenology of its main host plant (Bopp & Gottsberger 2004). Female moths pollinate female plants and oviposit on flower ovaries. After hatching, first instar larvae mine into the developing capsule to feed on maturing seeds, while later instars move to nearby capsules before pupating in the soil. Oviposition and larval development of *H. bicruris* occur primarily on *S. latifolia* (Bopp & Gottsberger 2004). Early studies emphasized the role of *H. bicruris* as a seed predator (Bopp 2003), whereas later studies demonstrated that the species is simultaneously an effective pollinator, with pollination by male moths, together with high egg and larval mortality and the abortion of infested fruits, substantially offsetting the costs of seed consumption (Jolivet & Bernasconi 2006; Labouche & Bernasconi 2010; Villacañas de Castro & Hoffmeister 2020). How environmental conditions shift the net outcome of this interaction along the mutualism– antagonism continuum remains largely unexplored. Plant chemistry is likely to play a central role in mediating these shifts. Several VOCs in the floral scent of *S. latifolia* trigger behavioural responses in *H. bicruris* (Dötterl et al. 2006) and attraction of female moths to female flowers has recently been shown to correlate with the abundance of a previously undescribed dimethylxanthine derivative and a previously undescribed pyrrolidine derivative in nectar (Zill *et al*. 2025), compound classes with known neuroactive or pollination-beneficial effects in other plant species (Wright *et al*. 2013).

### 2.2 Plant and pollinator cultivation

Seed capsules were collected in each of five European *S. latifolia* populations (***Figure 1b***). Plants were grown from seeds for one generation of controlled outcrossing within populations in the greenhouse (see Schrieber *et al*. 2017 for details). The resulting F1 generation was raised and prepared for the experiment as described in ***Note S1***.

To standardize pollinator background across plant populations and avoid confounding effects of local plant–pollinator matching, all experiments were conducted using a common laboratory population of *H. bicruris* from northern Germany, outside the geographic range of the experimental plant populations (***Figure 1b***). This design isolates plant population responses but does not address local plant–pollinator co-adaptation. A detailed description of the rearing conditions is provided in ***Note S1*.**

### 2.3 Experimental setup

We exposed female and male *S. latifolia* plants from all populations to well-watered and water-limited conditions in a fully factorial design (5 populations × 2 sexes × 2 watering regimes × 5 replicates = 100 plants, ***Figure 1c***). Relative soil moisture was maintained at ≥ 50% (well-watered) or ≥ 15% (water-limited), with 15% corresponding to the threshold at which all plants showed at least mild wilting in a pilot experiment. Soil moisture was measured every second day using a sensor (ML3 ThetaProbe Soil Moisture Sensor, Delta-T Devices Ltd., Cambridge, UK), and 450 mL water was added whenever moisture fell below the respective threshold to allow recovery. By enforcing the same external soil-moisture constraint relative to each plant’s demand, this approach standardises water limitation across sexes and populations that differ considerably in morphology. This treatment was applied for a total period of 12 weeks. Based on this setup, we quantified a comprehensive set of visual and chemical flower traits along with pollinator choice behaviour (***Figure 1d***, ***chapter 2.4)***, as well as quantitative and chemical seed traits along with *H. bicruris* larval performance (***Figure 1e***, ***chapters 2.5)***.

### 2.4 Data acquisition for floral traits and moth behaviour

#### 2.4.1 Visual flower traits

About four weeks after the onset of the watering treatment (two weeks after first hand pollination, see ***chapter 2.5***), we measured inflorescence height and counted the number of open flowers. Moreover, we selected one fresh but fully developed open flower (females: long papillous stigmata; males: dehisced anthers) per plant for the quantification of flower area and colour. A detailed description of these analyses is provided in ***Note S1***.

#### 2.4.2 Floral scent metabolome

About five weeks after the onset of the watering treatment (three weeks after first hand pollination, see ***chapter 2.5***) we started with the assessment of floral chemistry. Flower scent was characterised by trapping headspace VOCs on absorbent polydimethylsiloxane (PDMS) tubing following Kallenbach *et al*. (2014, 2015). Collection units were polypropylene cups with lids (50 mL Premium Line, Tedeco-Gizeh, Offenburg, DE) with 15 mm holes to prevent heat and waterlogging. Thirty minutes before the climate-chamber night cycle, one fully developed, open flower was inserted into each unit. After 1 h recovery, two PDMS tubes (5 mm length, Carl Roth, Karlsruhe, DE, solvent-and heat-cleaned) were threaded onto copper wire and inserted inside the unit. Tubes were removed after 6 h and stored at -18 °C in sealed glass vials. To minimise cross-contamination, five randomly selected plants were transferred to otherwise empty climate chambers and arranged at maximum spacing for VOC collection. A detailed description of sample analyses via thermal desorption gas chromatography mass spectrometry (TD-GC-MS) and compound annotation is provided in ***Note S1***. We detected a total of 45 VOCs in the floral headspace of *S. latifolia*. Of these 43 could be putatively identified via matches of RIs and mass spectra with databases. The majority of these VOCs have been previously described in *S. latifolia* and 19 of them have been shown to elicit antennal and/or behavioural responses in *H. bicruris* (Dötterl *et al*. 2006) ***(Table S1a)***.

#### 2.4.3 Floral nectar metabolome

Floral nectar was sampled from three fully developed, open flowers per plant. Nectar was extracted using 1 µL microcapillary tubes (Minicaps NA-HEP, Hirschmann Laborgeräte, Eberstadt, DE) from the ovary base in females and from the filament base and anthophore in males. Total nectar volume per flower was quantified, and the mean volume per plant was calculated. For metabolome profiling, capillaries were rinsed with 85 µL ultra-pure water (ROTISOLV Ultra LC-MS grade, Carl Roth, Karlsruhe, DE) into reaction tubes, pooling up to 6 µL nectar per individual (max. 2 µL per flower). Samples were vortexed, snap-frozen in liquid nitrogen, and stored at **−**80 °C. A nectar solution volume of 0.5 mL was mixed with 0.5 mL methanol and 4 mL methyl tert-butylether (both ROTISOLV Ultra LC-MS grade, Carl Roth, Karlsruhe, DE). After incubation (30 min, 25 rpm), phase separation was induced by adding 0.5 mL ultra-pure water. After further incubation (10 min, 25 rpm), and centrifugation (10 min, 4000 × g), the upper lipophilic phase was collected. The lower phase was re-extracted twice with methanol to recover hydrophilic metabolites. Hydrophilic and lipophilic extracts were dried at 0.1 bar and 45 °C (SpeedVac, Thermo Scientific, Germany) and resuspended in 0.5 mL of either water/methanol (50:50, v/v; hydrophilic) or isopropanol/chloroform (3:1, v/v; lipophilic). Prior to UHPLC-qTOF-MS/MS analysis, extract dilution was adjusted for differences in nectar volume among individuals, thereby standardizing the amount of original nectar represented in the analysed extract and preventing differences in sampled nectar volume from generating systematic dilution effects. A detailed description of sample analyses via UHPLC-ESI-qToF-MS/MS and compound annotation is provided in ***Note S1***. The untargeted analyses of hydrophilic and lipophilic extracts from a standardized nectar volume per plant yielded abundance data for ≈ 5,500 features. Of these, ≈ 120 features were tentatively identified via matches of their mass spectra with databases or via cheminformatics ***(Table S1b)***.

#### 2.4.4 Pollinator choice assays

Behavioural assays with *H. bicruris* were performed on an independent set of plants grown and treated under identical conditions because a disease outbreak temporarily left no adult moths available during flowering of the original plants. Consequently, only two of the five study populations, representing the greatest geographic contrast, were included. All other measurements reported in this study were conducted on the original experimental plants.

Behavioural assays took place under laboratory conditions between 8:00 pm and 0:30 am - the peak activity time of our captive moths. For the choice assays, female and male moths were kept individually in ventilated, transparent boxes (15 × 12 × 11 cm) equipped with shelters and a feeding station and were repeatedly used throughout the experiment. Prior to individual housing, moths had been maintained together ***(chapter 2.2)***, allowing mating to occur, and consequently differed in mating history and age at the time of the assays. Thus, rather than representing age- or mating-standardized individuals, the experimental moths encompassed natural variation in adult state likely to be encountered in the field. For each assay, the honey-water solution was removed in the morning and replaced in the evening with two fully developed, freshly picked flowers from the experimental plants, which were fixed in lidless Eppendorf tubes filled with tap water. The two flowers were positioned on opposite sides of the boxes for maximum spatial separation. Moths were provided with the binary choice among the following four treatment groups: control females versus water-limited females, control males versus water-limited males, control females versus control males, water-limited females versus water-limited males. Across both plant populations, 112 choice trials were conducted, with 24–30 replicate trials for each of the four choice combinations. These trials involved 28 moth individuals (10 females and 18 males), which were randomly assigned across the different choice trials. Behavioural observations were restricted to the first four flower visits of each moth, with a visit defined as starting and ending with physical contact with a flower. We deliberately decided against first-visit choice tests, as these give chance undue weight and moth individuals frequently revisit individual flowers or probe several flowers before feeding extensively or ovipositing on one under natural conditions (personal observation). Within this observation period, two behavioural parameters were quantified for each of the two flowers: (i) the number of feeding attempts, with an attempt starting and ending with unrolling and rolling up the proboscis and its insertion into and removal from the corolla tube, respectively; and (ii) the number of oviposition attempts.

### 2.5 Data acquisition for seed traits and larval performance

To disentangle maternal effects of water limitation on seed provisioning and their consequences for larval performance from indirect effects mediated by treatment-dependent pollinator visitation and reproductive investment, we standardised pollen receipt across treatments. From the second week of the watering treatment onward, female flowers were therefore hand-pollinated over three weeks, i.e. all flower and moth behavioural traits were acquired after pollination. Over two weeks, each female was pollinated with all five males from the same population and watering treatment, using pollen from two male flowers per female flower and equal numbers of female flowers per male. Pollen was applied by rubbing freshly dehisced anthers onto female stigmas. Pollinated flowers were covered with breathable mesh bags, and ripe capsules were harvested at week 12 of the experiment.

#### 2.5.1 Seed quantity and germination

For all female plants (5 populations × 2 watering treatments × 5 replicates = 50), we determined the number of flowers to which a pollination treatment was applied, total seed mass, total seed number, and individual seed area (latter two quantified using WinSeedle software, Regent Instruments, Quebec, CA). The following quantitative traits were considered for further analysis: *seed set per pollination attempt = total seed number / number of pollinated flowers, mean seed weight = total seed mass / total seed number, and mean seed* 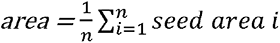.

Up to 50 seeds per female plant were placed on filter paper moistened with sterile water and set on a layer of 2 mm sterile glass beads in 90 mm Petri dishes. Dishes were incubated in a climate chamber (MLR-32, Panasonic Healthcare, Kyoto, JP) at 20 °C / 10 °C under a 16 h light / 8 h dark cycle. Germination was scored daily for 15 days and every third day for another 15 days. Germination was defined as seed coat rupture with visible radicle or cotyledons. Germination percentage, mean germination time, and synchronization index were calculated using GerminaR (Lozano-Isla *et al*. 2019).

#### 2.5.2 Seed metabolome

Fresh seeds were mortar-grind for 15 min with repeated addition of liquid N_2_. For extraction of polar and semi-polar metabolites, 10 mg seed powder was mixed with 0.8 mL pre-cooled methanol (-80°C) and 2 mL water (both ROTISOLV Ultra LC-MS grade, Carl Roth, Karlsruhe, DE), incubated for 10 min at 25 rpm on an end-over-end shaker, and centrifuged (4000 × g, 4 °C, 10 min). The supernatant was analysed via UHPLC-ESI-qToF-MS/MS as described for nectar. The methanol extract of *S. latifolia seeds* contained ≈ 3500 metabolic features of which ≈ 230 features were tentatively identified via matches of their mass spectra with databases or via cheminformatics ***(Table S1c)***.

#### 2.5.3 Larval bioassays

We assessed the effects of water limitation on seed consumption, growth, and survival of *H. bicruris* larvae. We selected 50 third instar larvae of comparable developmental stage (1.0 to 1.5 cm), as earlier life stages do not feed on ripe seeds. Larvae were weighed and assigned to pairs based on similarity in body mass. They were kept individually in ventilated plastic containers lined with filter paper and provided with an empty *S. latifolia* seed capsule as shelter. Each larva was fed seeds from a single female plant. Within each weight-matched pair, larvae were offered seeds from the same population × maternal family combination but from different watering treatments. Every four days, larvae were supplied with fresh seeds (crushed for all larvae < 2 cm and intact thereafter) and containers were cleaned of frass and seed leftovers. Seed supply always exceeded demand, and surplus seeds remained at each feeding. To avoid seed imbibition that could bias the assessment of remaining seed mass, larvae were transferred every other day for 2 h to an empty container where droplets of tap water were provided for drinking. Larvae were weighed throughout the experiment to determine the final body mass immediately prior to onset of the pre-pupal weight loss. Relative growth rate was calculated from the initial (t1) and this maximum pre-pupal (t2) body mass as: *RGR = [ln(weight t2) – ln(weight t1)]/ t2-t1*. Total seed consumption was calculated as the cumulative difference between the mass of seeds supplied and the mass remaining up to this time point. Time to pupation was quantified as the number of days from the start of the feeding experiment until its occurrence.

### 2.6 Statistical analyses

#### 2.6.1 Univariate traits

All univariate response traits were analysed in R 4.4.1 (R Development Core Team 2024) using (generalized) linear mixed effects models - (G)LMMs (R-package “glmmTMB”, Brooks *et al*. 2017). Models were structured as follows: Visual flower traits, nectar volume, quantitative seed traits, germination, and larval performance traits were modelled as responses. Fixed effects were watering treatment (factor: well-watered, water-limited; all models), plant sex (factor: female, male; excluded from seed trait and larval performance models), and 20-year mean Climate Moisture Index of the three hottest months (CMI, continuous: **−**123.05 – **−**50.99; included in all models as a covariate to account for climatic variation among population origins), as well as all possible interactions among these variables. Larval performance models additionally included consumed seed mass (relative growth rate) or initial larval age (time to pupation) as additional covariates (both continuous, additive and interactive effects). Plant population was set as random effect in all models.

Moth choice assays were analysed with two binomial models testing (i) preferences for a specific watering treatment within sexes (assays: control female *versus* water-limited female, control male *versus* water-limited male); and (ii) preferences for a specific plant sex within watering treatments (assays: control female *versus* control male, water-limited female *versus* water-limited male). Model i / ii included the proportion of feeding or oviposition attempts directed towards control relative to water-limited flowers (i) or towards female relative to male flowers (ii) as binomial response and flower sex (i) or watering treatment (ii) as predictors. Further predictors included in both models were moth sex (except for oviposition), plant population, and all possible interactions among predictors. Moth individual was set as random effect.

All models were evaluated for compliance with underlying assumptions using diagnostic tests and plots from the R-package "DHARMa" (Hartig 2022) and, where appropriate, were optimized with regard to error distribution, dispersion or zero-inflation. For downstream analyses, we calculated type III ANOVA tables as omnibus tests of fixed effects (based on Wald-χ² tests) while setting all contrasts for factors to sum-zero coding (R-package “car”, Fox & Weisberg 2019). If factorial predictors were involved in significant interactions, we additionally calculated simple effects (i.e., post-hoc tests) based on estimated marginal means. (R-package “emmeans”; Lenth, 2024). For moth choice models, post-hoc contrast (R-package “emmeans”; Lenth, 2024) were used to test for a significant deviation of the binary outcome from the intercept of 0.5 (i.e., preference or avoidance) within the different plant sex × moth sex × plant population (i) or watering treatment × moth sex × plant population (ii) combinations.

#### 2.6.2 Metabolome traits

Statistical analyses of metabolome data were identical for floral scent, nectar, and seeds. We performed permutational analysis of variance (PERMANOVA, R-package “vegan” Oksanen *et al*. 2024). The Bray Curtis dissimilarities of the full datasets (scent: n = 45 features; nectar: n = 5500 features; seed: n = 3560 features) were used as response distance matrix. Fixed effects were watering regime (factor: well-watered, water limited), plant sex (factor: female, male; excluded for seeds) and their interaction. Populations were set as strata (random effect equivalent). Population level CMI was not included as fixed effect because PERMANOVA cannot test covariates that are constant within a stratum, and because analyses based on population-level averaged metabolomes showed at most minor CMI effects ***(Note S2)***. The models ran with 100,000 permutations. Multivariate group separations were visualised using principal coordinates analysis (PCoA) based on Bray–Curtis distance matrices using multidimensional scaling. We verified that negative eigenvalues were negligible relative to the leading positive eigenvalues, indicating that the PCoA provided an adequate representation of the Bray–Curtis distance structure.

To identify the compounds responsible for multivariate group structure in a next step, we used an approach that is comparable to a volcano plot and simultaneously allows the statistical assessment of interaction effects among plant sex and watering treatment, as well as accounting for the highly heterogeneous distributional properties of GC/LC–MS metabolite intensities. Instead of applying two-group significance tests, we entered all metabolites as responses into a (G)LMM. This procedure followed a two-step logic: (1) identification of the best-fitting model and (2) downstream analysis of that model. A detailed description of this approach is provided in ***Note S1***.

## 3 Results

### 3.1 Effects of watering regime and plant sex on floral traits and pollinator behaviour

Visual floral traits were shaped primarily by the main effects of watering regime and plant sex, whereas their interaction had no effect (***Figure 2***, ***Table S2)***. Water limited plants produced significantly shorter inflorescences, and fewer flowers with smaller areas. In comparison to male plants, females produced significantly fewer flowers (after pollination only) with larger areas and lower green:blue reflectance ratios.

**Figure 2:**
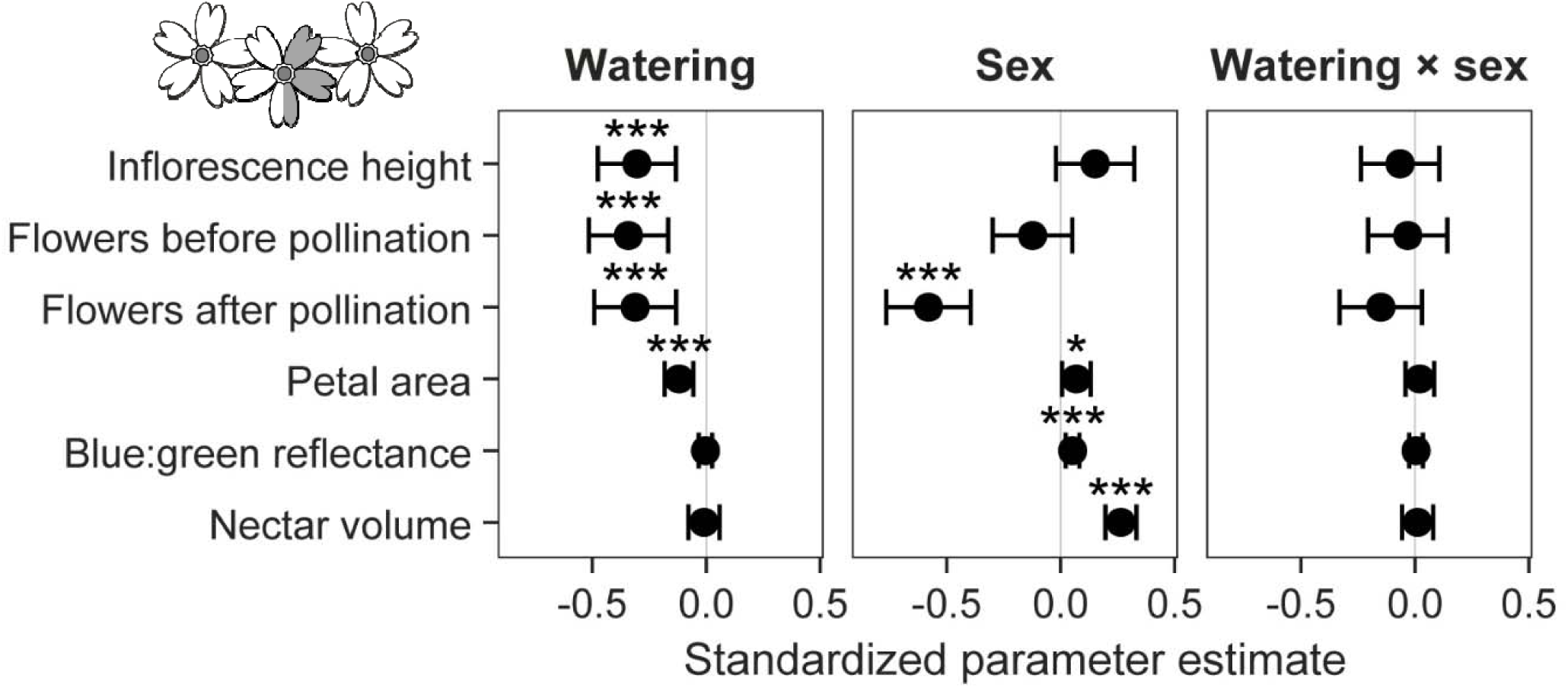
Forest plot summarizing the effects of watering regime, plant sex and their interaction on visual floral traits and nectar quantity of Silene latifolia. Standardized coefficients of (G)LMMs are given with 95% confidence intervals (bars) and significance levels assessed via type III ANOVA. Negative coefficients indicate lower trait values in water-limited compared to well-watered plants (watering), in male compared to female plants (sex), and weaker watering treatment effects in male compared to female plants (watering × sex). N = 5 populations × 2 sexes × 2 watering regimes × 5 replicates = 100.

The composition of floral scent assessed via PERMANOVA was likewise mainly shaped by the main effects of plant sex and watering regime ***(Figure S1a, Table S2)***. PCoA centroids segregated mainly between female and male plants, whereas water limitation primarily increased variation in scent composition rather than shifting centroids. The results of the model loop were consistent with these patterns, with several metabolites modulated clearly by the main effect of sex, but weak effects of watering regime, or the interaction sex × watering regime ***(Figure S1b–f, Table S1a)***. Male flowers showed a strong enrichment (p_FDR_ < 0.05, log_2_FC < -1) of the *H. bicruris* core attractants guaiacol and veratrole. Moreover, many other behaviourally active VOC including the *H. bicruris* core attractants Lilac aldehyde A-D had significantly (p_FDR_ < 0.05) but moderately (log_2_FC < -0.5) higher abundances in males.

The nectar volume per flower was not affected by watering regime, but markedly higher in females than males across both watering regimes (***Figure 2***, ***Table S2)***. The composition of the nectar metabolome assessed via PERMANOVA revealed a significant effect of the interaction watering regime × plant sex ***(Table S2)****. PCoA* centroid shifts in response to the watering regime were clearly weaker in females than in males and occurred along different ordination axes in both sexes (***Figure 3a***). The results of the model loop were consistent with these patterns: a comparably small fraction of metabolites was modulated by the main effects of watering regime and plant sex, while the largest fraction was modulated by their interaction (***Figure 3b***, ***Table S1b)***. In the following description of effect directions, numbers outside and inside parentheses refer to metabolites with *|log2FC| > 1* and *p_FDR_ < 0.05* or *p_RAW_ < 0.05*, respectively. Watering regime had a weak but consistent negative main effect with 3 (48) features decreasing under water limitation. These included a phytoecdysteroid, the antioxidants phylloquinone and 2,4,6-tri-tert-butylphenol, and four primary fatty acid amides (***Figure 3c***, ***Table S1b)***. The main effect of sex was more pronounced with 12 (42) features having higher abundances in females than males. These included the amino acid (derivatives) L-arginine, L-pyroglutamate, L-glutamine, and N²-acetyl-L-ornithine, two flavones, the volatile terpenoids D-camphor and carveol, and two additional non-volatile terpenoids. Finally, two metabolites that were first described in a previous study on *S. latifolia* nectar, where they were correlated with *H. bicruris* behaviour, were also included: a novel, structurally unresolved dimethylxanthine derivative and a novel, structurally unresolved pyrrolidine alkaloid (***Figure 3e***, ***Table S1b, Note S3)***. On the other hand, 10 (76) features had lower abundances in females than males. These included sucrose, a phytoecdysteroid, an unknown oligopeptide, and the antioxidants glutathione and phosphocholine. The interaction sex × watering treatment had the most pronounced effects on nectar composition. When focusing on differential responses to water limitation between sexes, two effect directions were clearly dominant (***Figure 3f***): A total of 1408 (781) features decreased in males but not in females. Of these we (putatively) identified three small peptides, numerous lipid-derived metabolites (e.g., a prostaglandin-like C20 oxylipin), three phenylpropanoids (e.g., hydrocinnamic acid), two nitrogen-containing compounds (an aromatic carbamate and a piperidine derivative) and two terpenoids (e.g., a precursor of the floral VOC linalool). In contrast, only 0 (6) features decreased in females but not in males (e.g., a putative carotenoid and small peptide). Given the standardization of analysed nectar amounts, these sex- and compound-class-specific reductions (lipids) are unlikely to reflect systematic differences in sample dilution. At the same time, 3 (1) features increased under water limitation only in males (all remained unannotated), whereas 6 (5) features increased only in females (e.g., a phosphocholine and a triterpene). When focusing on sexual differences within watering treatments, corresponding dominant effect directions emerged (***Figure 3f***): Males had higher abundances of numerous metabolic features under well-watered conditions, whereas water limitation largely eliminated this sexual dimorphism and, for some features, even reversed its direction.

**Figure 3:**
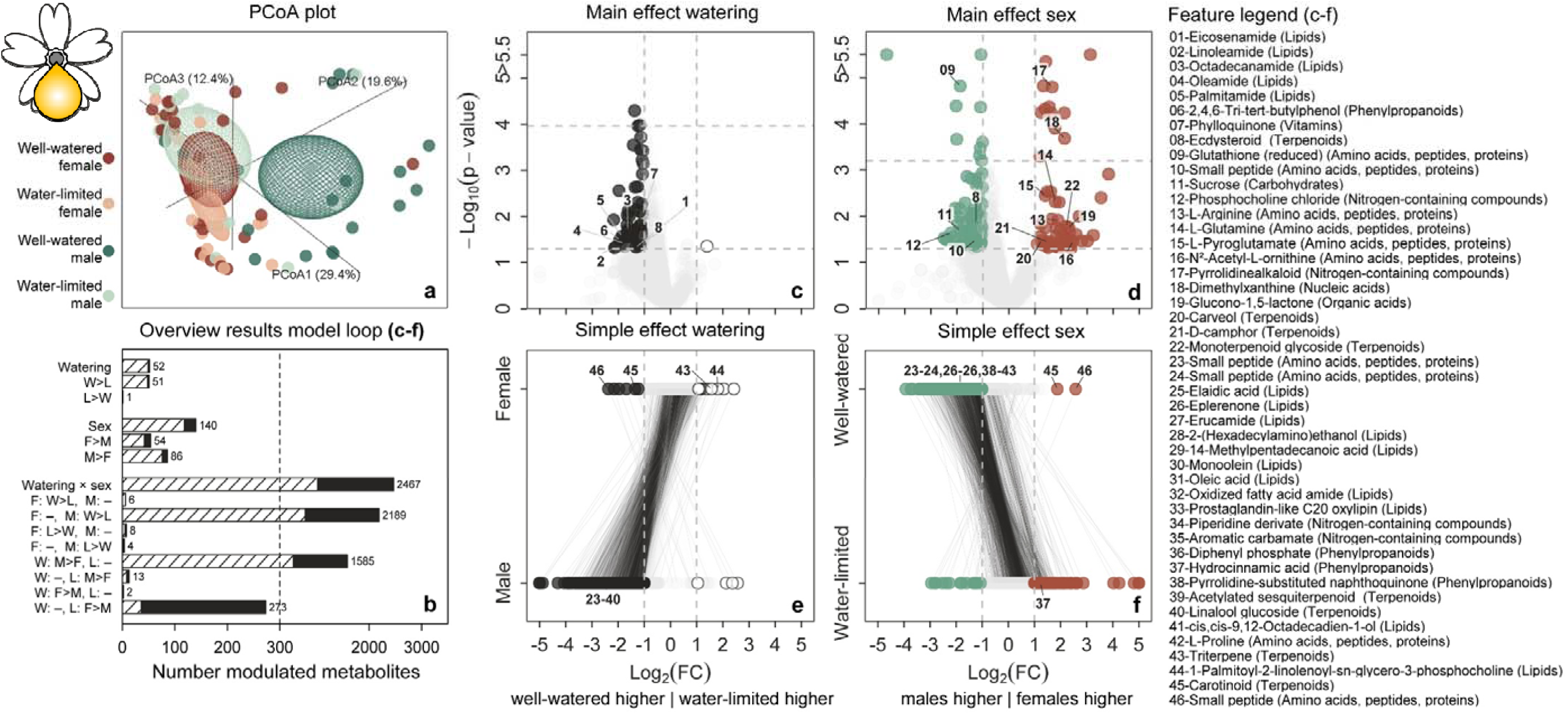
Effects of watering regime and plant sex on nectar metabolome composition in Silene latifolia. (a) 3D PCoA plot with points representing well-watered females (W–F, dark red), water-limited females (L–F, light red), well-watered males (W–M, dark turquoise), and water-limited males (L–M, light turquoise). Ellipsoids represent 95% confidence regions around group centroids. (b) Bar plot summarizing mixed model-loop results as the number of features modulated by each effect and direction (|log2FC| > 1 and p / pFDR <0.05 [dashed / filled]). (c,d) Volcano plots showing the main effects of watering regime and plant sex for all features not additionally involved in a significant interaction. Modulated features (p / p_FDR_ <0.05 [lower / upper horizontal dashed line] and log_2_FC < -1 / > 1 [left / right vertical dashed line]) are coloured: higher in well-watered plants= dark grey, higher in water-limited plants = white, higher in females = red, higher in males = turquoise. Interaction effect volcano plots show (e) the simple effect of watering regime within females and males, and (f) the simple effect of plant sex within well-watered and water-limited plants. Identical fea ures within these group comparisons are connected by lines. For further details on the identity of features, see Table S1b. N = 5 populations × 2 sexes × 2 watering regimes × 3–5 replicates = 80.

*Hadena bicruris* choice of well-watered relative to water-limited plants in terms of feeding attempts depended on the interaction moth sex × plant population (***Figure 4a***, ***Table S3)***, with only female moths showing a significant preference for flowers from well-watered plants in one of the two tested populations (Pop 2 = lowest CMI). For oviposition, moths generally preferred flowers from well-watered over water-limited plants (***Figure 4b***, ***Table S3)***. Similarly, moth choice between female and male flowers in terms of feeding attempts depended on plant population, with a significant preference for male flowers restricted to the same population as above (***Figure 4c***, ***Table S3)***. Finally, female moth preference for ovipositing on female relative to male plants depended on watering treatment, with a significant preference for female flowers under well-watered but not water-limited conditions (***Figure 4d***, ***Table S3)***.

**Figure 4:**
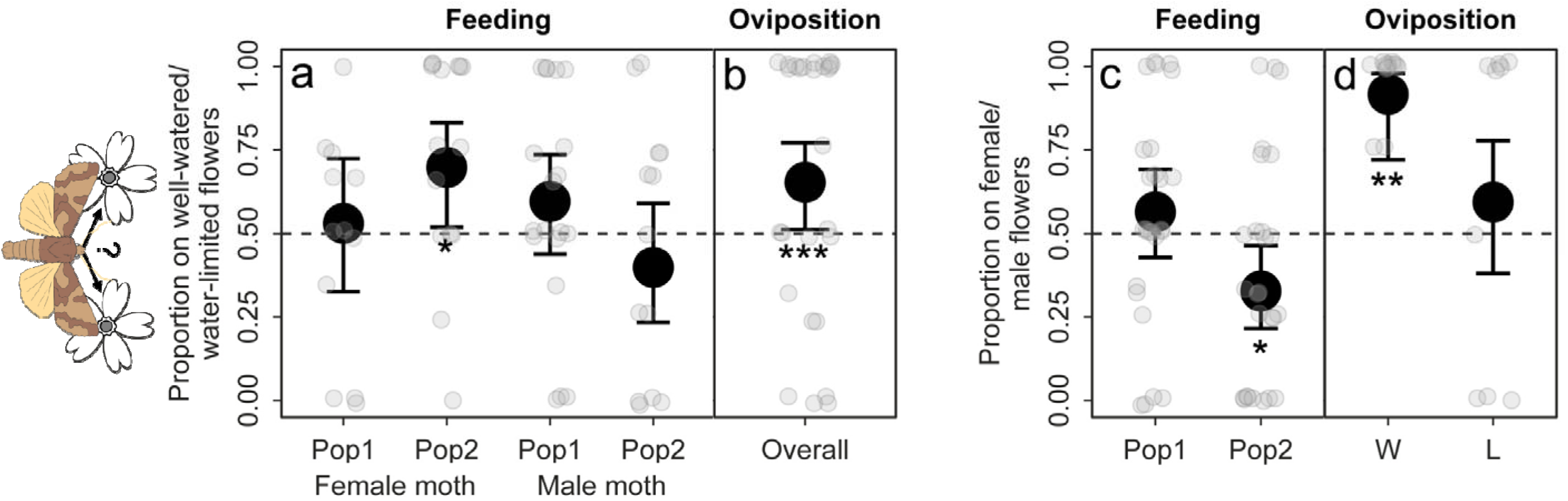
Effects of watering regime, plant sex, plant population and moth sex on choice behaviour of Hadena bicruris. Large circles represent estimated marginal means from the GLMMs with 95% confidence intervals, while smaller jittered circles show observed proportions. (a, b) Choice of flowers from well-watered relative to water-limited plants, with values > 0.5 indicating preference for well-watered plants. Feeding preference depended on plant population and moth sex (a), whereas oviposition showed an overall preference for well-watered plants (b). (c, d) Choice of female relative to male flowers, with values > 0.5 indicating preference for female flowers. Feeding preference depended on plant population (c), whereas oviposition preference depended on watering regime (d). N = 112 choice trials across 4 choice combinations (24–30 replicates each), using 28 repeatedly tested moths (10 females, 18 males); two choice combinations were pooled per analysis.

### 3.2 Effects of watering regime on plant seed traits and larval performance

Seed set per pollination attempt and seed weight:area ratio were significantly reduced under water limitation (***Figure 5***, ***Table S4)***. Germination responses were unaffected ***(Table S4)***.

**Figure 5:**
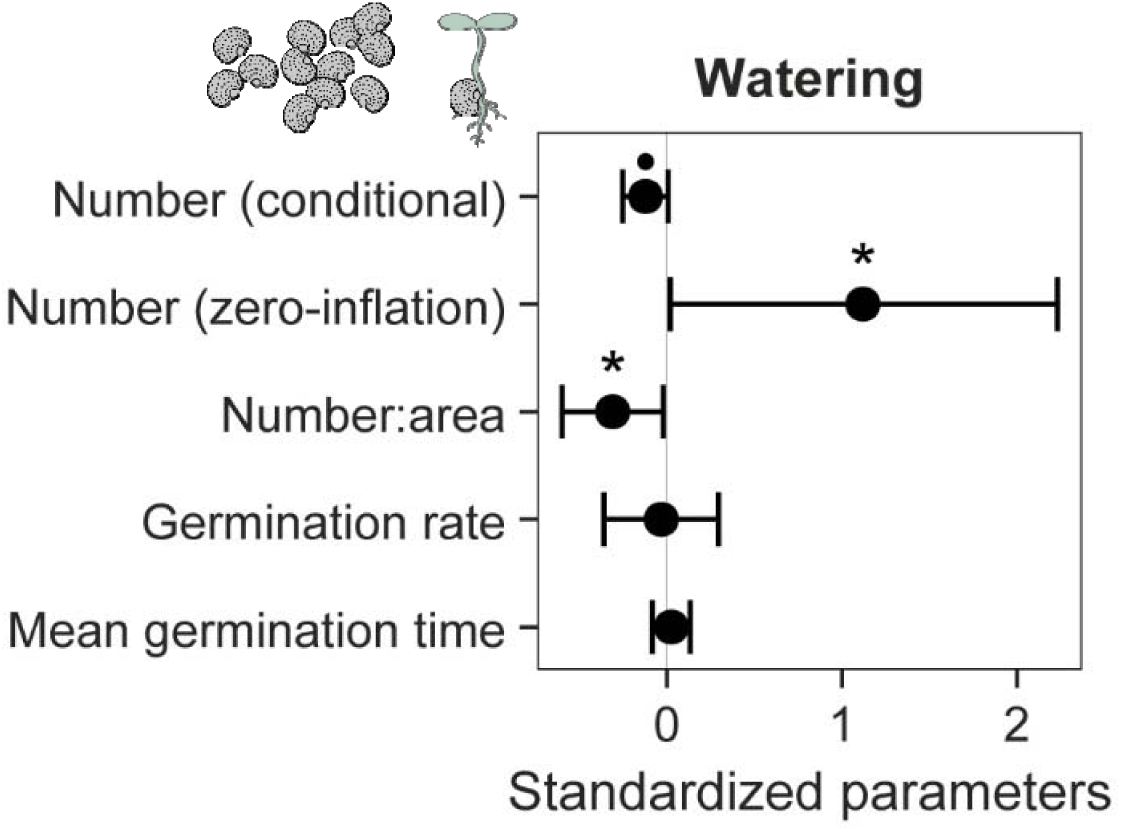
Forest plot summarizing the effects of watering regime on seed traits of Silene latifolia. Standardized coefficients of (G)LMMs are given with 95% confidence intervals (bars) and significance levels assessed via type III ANOVA. Negative coefficients indicate lower trait values in water-limited compared to well-watered plants (watering). N = 5 populations × 1 sex (female) × 2 watering regimes × 5 replicates = 50.

The overall metabolic composition of *S. latifolia* seeds assessed via PERMANOVA and PCoA was largely unaffected by watering regime (***Figure 6a***, ***Table S4)***. However, subsequent model-loop analyses (***Figure 6b***, ***Table S1c)*** identified a weak yet consistent water limitation signature, characterized by reduced abundance of 35 (178) metabolites and increased abundance of 15 (144) metabolites with numbers outside and inside parentheses referring to features with *pRAW < 0.05* and *|log2FC| > 1*, or *pRAW < 0.05* only, respectively. Annotated reduced metabolites included several small peptides, numerous carbohydrates (oligosaccharides such as melibiose, scyllo-inositol, N-Acetyl-galactosamine, N-Acetyl-mannosamine), nucleic acids (cytosine, guanosine, cytidine, xanthine, xanthosine) and several putative defense compounds (a putative iridoid glycoside, two putative benzoxazinoids, a coumarin and a nicotinic acid derivative). In addition, the previously described novel dimethylxanthine derivative ***(Note S3)*** also occurred in seeds and showed a marginally significant decrease under water limitation. Annotated increased compounds mainly included lipids (Eicosadienoyl ethanolamine; hydroxystearic acid, pentadecatetraenoic acid, erucamide, a prostaglandin-like C20 oxylipin), a disaccharide polyol, and adenosine 5’-monophosphate.

**Figure 6:**
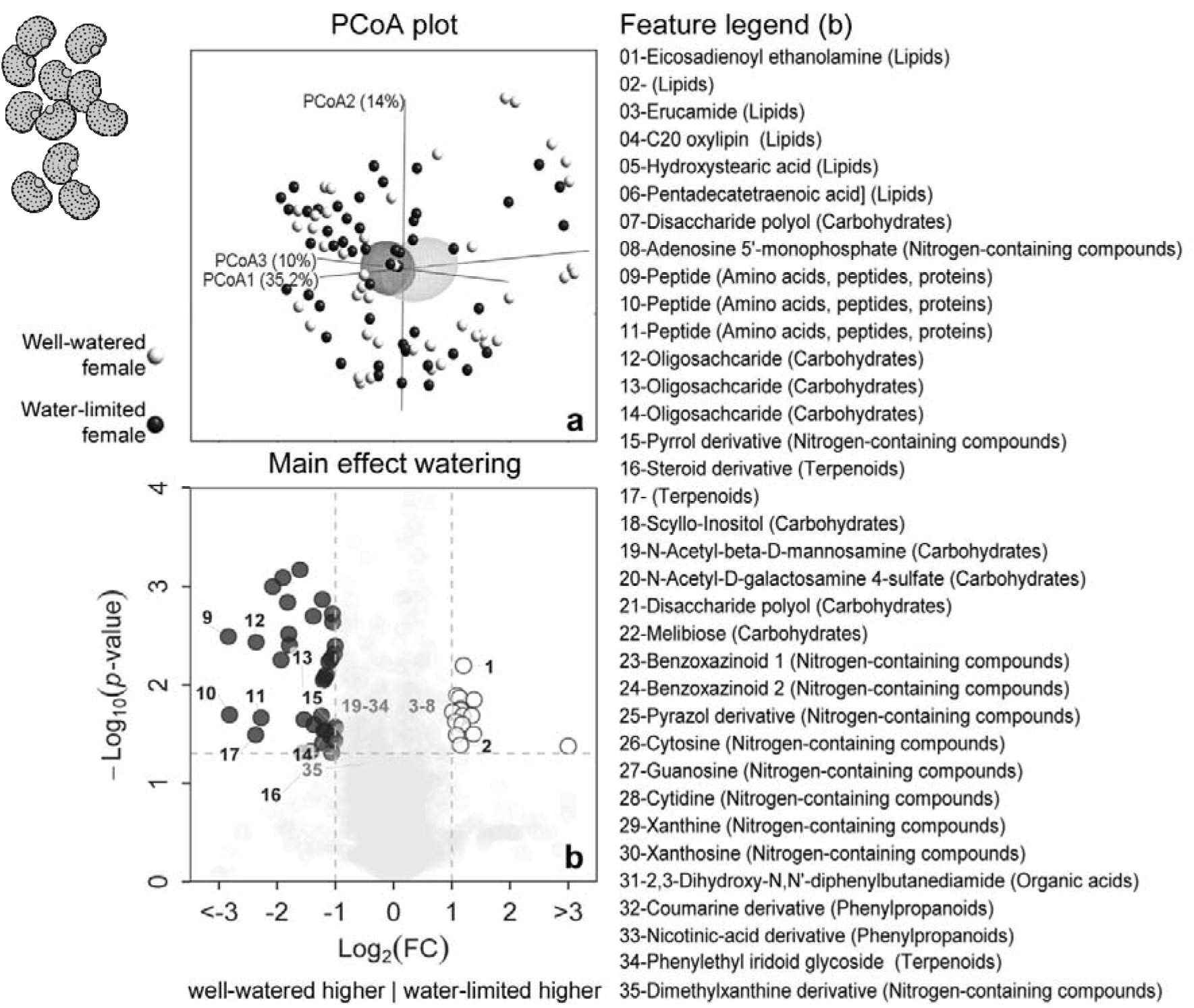
Effects of watering regime on seed metabolome composition in Silene latifolia. (a) 3D PCoA plot with points representing well-watered female plants (black), and water-limited female plants (white). Ellipsoids represent 95% confidence regions around group centroids. (b) Volcano plot showing the results of the model loop for the main effects of watering regime. Modulated features (p_RAW_ <0.05 [horizontal dashed line] and log_2_FC < -1 / > 1 [left / right vertical dashed line]) are coloured: higher in well-watered plants= black, higher in water-limited plants = white. For further details on the identity of features, see Table S1c. N = 5 populations × 1 sex (female) × 2 watering regimes × 3–5 replicates = 40.

The relative growth rate of *H. bicruris* larvae depended on the two-way interaction watering treatment × consumed seed mass (***Figure 7a***, ***Table S4)***. Growth rates increased significantly with the consumed mass of seeds from water-limited plants only (*water-limited: partial slope = 0.032, SE = 0.006, p < 0.001; well-watered: partial slope = −0.002, SE = 0.007, p = 0.731*). Consistent with this, time to pupation was shorter when larvae fed on seeds of water-limited as compared to well-watered plants (***Figure 7b***, ***Table S4)***. Pupation rate and the total seed mass consumed during larval development were not affected by watering regime and covariates.

**Figure 7:**
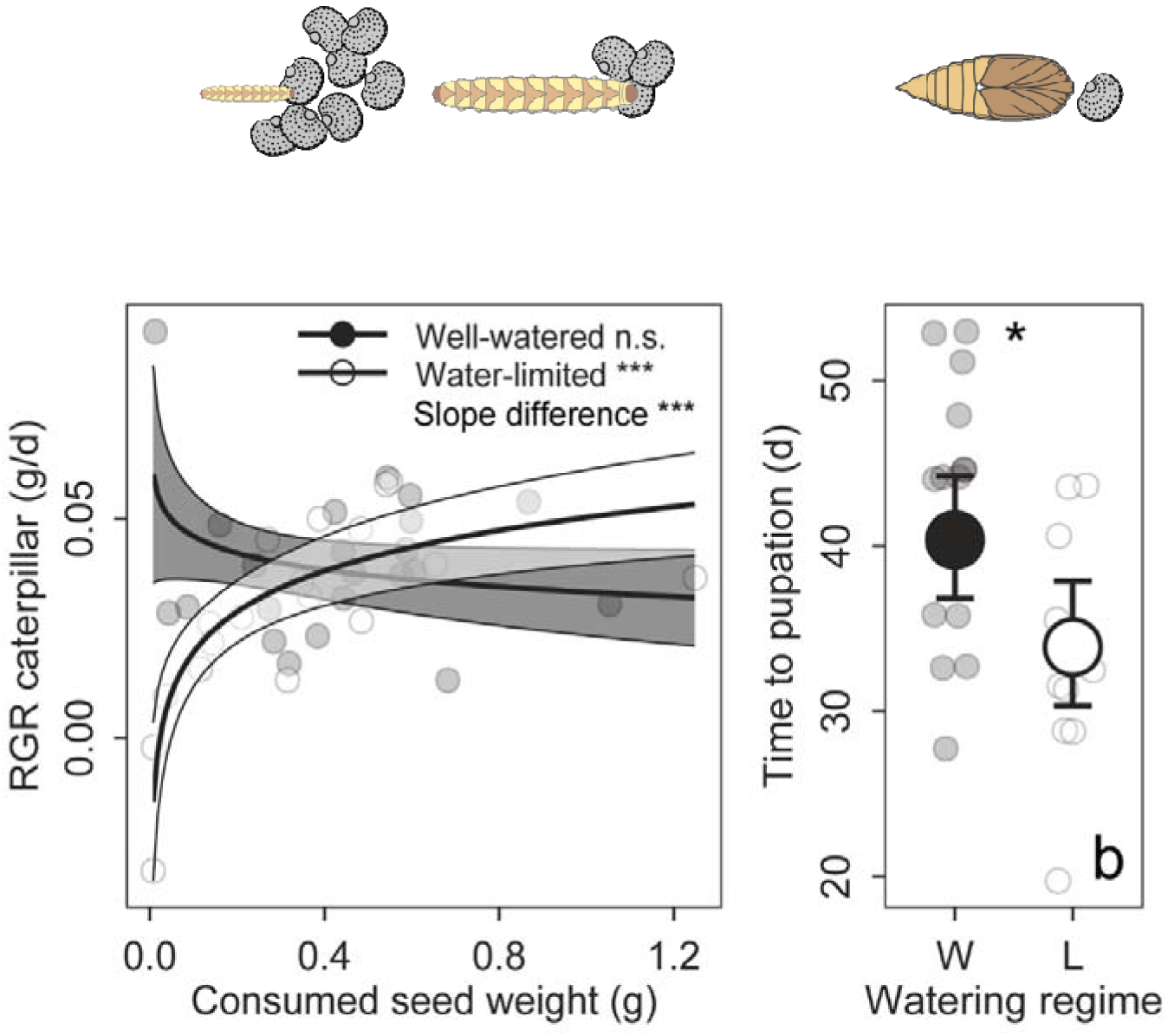
Effects of watering regime on the performance of Hadena bicruris larvae feeding on seeds from well-watered (W = black) and water-limited (L = white) plants. (a) Regression plot illustrating the relative growth rate of larvae in dependence of the seed weight consumed from W and L plants. Lines represent the prediction of the GLMM with 95% CI (shades). (b) Effects of watering regime on the time to pupation. Large circles represent marginal estimated means from the GLMM with 95 % CI (bars), while jittered smaller circles represent observed data. N = 5 populations × 1 sex (female) × 2 watering regimes × 3–5 replicates = 40.

## 4 Discussion

### 4.1 Water limitation reshapes multidimensional floral phenotypes and their sexual dimorphism, with consequences for a nursery pollinator

Water limitation clearly compromised visual floral traits integrating resource availability over longer developmental periods in both female and male *S. latifolia* (inflorescence height, flower number, petal area; ***Figure 2a***). In contrast, floral scent emission, often reported to be sensitive to water limitation owing to its integration with ABA stress-response pathways (Jaworski *et al*. 2022; Ré *et al*. 2011), was only weakly constrained ***(Figure S1c)***. Even more strikingly, floral traits directly linked to plant water status, such as colour composition through cell turgor and nectar volume, remained unaffected (***Figure 2a***). Likewise, nectar sugar and amino acid abundances remained stable (***Figure 3c***), indicating maintenance of the main nutritional rewards in both sexes. Together, these responses suggest a shift in resource allocation under water limitation, with investment in whole-plant floral display being reduced while maintaining the attractiveness and reward quality of fewer individual flowers. Beyond these main nutritional components, water limitation induced substantial shifts in nectar chemistry in both sexes, with most responsive metabolites decreasing in abundance and primary fatty acid amides (FAAs) particularly prominent (***Figure 3c***). FAAs are phylogenetically widespread yet understudied constituents of floral nectar (Vannette 2026). In plants, primary FAAs are excreted into the rhizosphere, where they modulate microbial interactions (Miao *et al*. 2025; Sun *et al*. 2016), making similar functions in nectar plausible. As insects possess FAA hydrolases, primary FAAs may serve as metabolic fuel for obligate nectarivores such as moths, as shown for fatty acids (Kim *et al*. 2010; Levin *et al*. 2017). Moreover, the FAA oleamide may modulate behaviour, as it exerts neurobiological effects involving the endocannabinoid system in mammals, while endocannabinoid-like lipids affect food preference and consumption in insects (He *et al*. 2021; Mendelson & Basile 2001).

Alongside the main effects of water limitation, pronounced main effects of plant sex were evident across floral traits. Previously described sexual dimorphisms in visual traits, floral scent, nectar volume, and sugar abundance were largely confirmed here (***Figure 2b***, ***Figure S1e,*** ***Figure 3e***) and have been discussed extensively in an evolutionary and functional context elsewhere (e.g., Waelti *et al*. 2009). More importantly, we identified previously undescribed constitutive sexual dimorphisms in nectar metabolite composition. Female nectar contained higher abundances of the amino acids L-arginine, L-glutamine, and L-pyroglutamate (likely a product of ESI-induced cyclisation of L-glutamate). L-Arginine provides nitrogen and precursors for nitric oxide and polyamines involved in insect reproduction and oogenesis (Smykal & Raikhel 2015). L-glutamine and L-glutamate are considered central to nectarivore nutrition and are known to exert phagostimulatory effects (Nicolson 2022). Females also had higher abundances of terpenoids (e.g., carveol, D-camphor) that may contribute nectar taste and perception by floral visitors (Barberis *et al*. 2023). Finally, females had higher abundances of a novel pyrrolidine alkaloid and novel dimethylxanthine derivative previously described by Zill et al. (2025), both of which positively correlated with the attraction of female moths in this previous study. Together, these patterns indicate that female nectar may specifically promote interactions with female moths through compounds associated with both their attraction and reproductive physiology, contrasting with the expectation that female plants should primarily evolve traits that deter their seed-predating pollinators (Burkhardt *et al*. 2012). Males, on the other hand, had higher abundances of sucrose – one of the main nutritional carbohydrates in nectar, reduced glutathione – a major antioxidant, phosphocholine – a potential source of choline and phosphate for insect nutrition, and a phytoecdysteroid – a plant steroid capable of modulating insect endocrine signalling and potentially affecting insect reproduction (Das *et al*. 2021). In summary, this extensive sexual dimorphism in nectar chemistry may promote pollinator movements between the sexes through complementary nutritional rewards, behavioural modulators, and physiological regulators, thereby facilitating pollen transfer in this dioecious system.

Beyond the main effects of water limitation and plant sex, nectar chemistry also showed pronounced interactive responses. These interactions were driven by a group of metabolites that were more abundant in male than female nectar under control conditions, but for which this sexual dimorphism disappeared or even reversed under water limitation because only males exhibited marked reductions in abundance. Lipids were again prominently represented among these reduced compounds, spanning a broad range of classes, including fatty acids, fatty acid amides, oxylipins, membrane lipids, and glycerolipids. As discussed above, these metabolites may contribute to microbial interactions, pollinator nutrition, and behavioural modulation, while membrane-active lipids may additionally influence nectar surface tension. The stronger reduction of lipid components in male nectar may reflect resource reallocation away from carbon- and energy-demanding constituents under stress. This may be particularly relevant in males because, despite producing less nectar per flower, their substantially greater flower production (***Figure 2***) and turnover likely require higher overall investment in nectar provisioning. Ultimately, the ecological significance of these male-specific metabolic shifts depends on whether they translate into altered pollinator behaviour.

Indeed, the behavioural responses of *H. bicruris* partially correlated with water- and sex-specific changes in multidimensional floral phenotypes (***Figure 4***). Feeding preferences were highly population-specific, both in response to water limitation and plant sex. Preferences for well-watered over water-limited flowers and for male over female flowers were each restricted to one of the two tested plant populations, notably the same population in both cases. This population specificity illustrates that both responses to environmentally induced changes and the expression of sexual dimorphism in floral phenotypes are highly context-dependent, which points to locally divergent selective pressures. Oviposition responses were considerably more consistent. Female moths generally preferred well-watered over water-limited flowers and preferred female over male flowers under well-watered conditions. However, the latter preference disappeared under water limitation. This closely parallels the sex × watering effect on nectar chemistry, where pronounced sexual dimorphism under well-watered conditions was strongly eroded by water limitation. Together, these patterns provide first correlative evidence that female moths may use sexually dimorphic nectar chemistry to discriminate between plant sexes and that water limitation interferes with this discrimination. Such impaired discrimination may be particularly costly for female moths because successful larval development depends on oviposition on seed-producing female plants. Direct manipulation of nectar chemistry will be required to isolate its contribution from other floral traits. In general, behavioural assays address immediate attraction and thus capture only one potential pathway through which changes in nectar chemistry under water limitation may affect *H. bicruris*. Allowing moths to complete their entire life cycle on plants will therefore be essential to determine whether such immediate behavioural responses scale up to consequences for individual fitness.

### 4.2 Water limitation reshapes seed chemistry, with consequences for larval performance in a nursery pollination system

In nursery pollination systems, seeds represent both the plant’s reproductive investment and the food source of developing insect larvae. We show how water limitation modifies this shared resource, with consequences for both interaction partners.

Seed set in hand-pollinated *S. latifolia* females was reduced under water limitation, notably as a qualitative rather than quantitative response, with ≈ 30 % of water-limited females producing no seeds at all (***Figure 5***). Seed set failure may arise from the individual or combined effects of poor male pollen quality, direct negative drought effects on female physiology (i.e. stress), and an adaptive female response to ensure survival and reproduce in more favourable years (Barnabás *et al*. 2008). Among females that successfully produced seeds, seed mass per area was significantly reduced under water limitation, a change typically associated with faster imbibition and germination as well as reduced filling of storage or seed coat tissues (Fenner 2022). Our results do not support an effect on germination (***Figure 5***), whereas seed metabolome analyses partially provide support for the latter interpretation.

Our analyses revealed both metabolite increases and decreases under water limitation (***Figure 6***). Increased metabolites comprised lipids and lipid-derived metabolites such as fatty acid amides, hydroxylated fatty acids and oxylipins. These compounds are commonly associated with plant drought stress signalling, membrane protection and plant defence against microbes, but may also serve as nutrient sources for germinating seedlings (Kaur *et al*. 2021; Kuźniak & Gajewska 2024; Sharma *et al*. 2023). Decreased metabolites included several small peptides, numerous carbohydrates, and nucleic acids (***Figure 6***). Some of these well-known seed storage compounds exhibited comparatively high relative abundances in the metabolome, and their reduction is therefore likely to have contributed to reduced seed filling (Bewley & Black 1994). Whether resulting from direct stress effects or resource reallocation towards stress resistance during seed provisioning, these changes are likely to reduce energy availability and the capacity for protein and nucleic acid synthesis during early seedling establishment despite unaffected germination (Barnabás *et al*. 2008). Moreover, many defence-associated compounds were also reduced under water limitation (***Figure 6***). These included an iridoid glycoside, a coumarin and a p-coumaroyl conjugate, benzoxazinoids, and the novel dimethylxanthine derivative (marginal effect only) as well as its potential precursor xanthosine ***(Note S3)***. Together, these findings suggest that maternal water limitation may propagate through reduced chemical defence of dormant seeds and seedlings, potentially increasing their susceptibility to pathogens, predators and herbivores.

Indeed, our study provides evidence that these changes in seed chemistry affected the physiology of *H. bicruris* larvae. Larvae fed seeds from water-limited plants grew increasingly faster with increasing seed consumption and reached pupation earlier than larvae fed control seeds, despite consuming similar amounts of seed tissue (***Figure 7***, ***Table S4)***. Although larvae may have profited from the increased abundance of energy-rich lipids (e.g., hydroxystearic acid), a simultaneous reduction of carbohydrates, peptides and nucleic acids argues against a generally improved nutritional quality. Instead, the concurrent reduction of multiple qualitative defences may have facilitated this more efficient conversion of consumed seed material into larval growth, as all of the above-mentioned substance classes are known to interfere with the performance of juvenile Lepidoptera (Wink 2010; Wouters *et al*. 2016). Resistance against specialist herbivores may in fact emerge only from the combined effects of multiple qualitative defences rather than from any single compound (Petrén *et al*. 2024; Richards *et al*. 2016). In *S. latifolia*, mechanisms limiting the costs imposed by *H. bicruris* have so far mainly been attributed to altered floral scent composition following pollination, selective fruit abortion, and mechanical barriers such as thick fruit walls (Burkhardt *et al*. 2009; Jolivet & Bernasconi 2006; Muhlemann *et al*. 2006). Our study provides the first evidence that a diverse mixture of defence compounds may contribute to limiting, rather than eliminating, the pollinating seed predator. Individual and combinatorial bioassays using artificial diets supplemented with naturally occurring concentrations of these compounds are a next step towards elucidating their function and environmental determinants.

### 4.3 Methylxanthine metabolism in *Silene* as a potential link between adult host choice and larval performance

In this study, we detected a dimethylxanthine derivative in both nectar and seeds of *S. latifolia*. Complementary NMR analyses revealed xanthosine in seeds and methylxanthine-related signals in seedlings, while genome annotation identified putative methyltransferases potentially involved in methylxanthine biosynthesis ***(Note S3)***. Together, these findings provide convergent evidence for methylxanthine metabolism in *S. latifolia*. Methylxanthines are neuroactive alkaloids that can influence insect behaviour and feeding, potentially contributing to pollinator interactions and plant defence (Stevenson *et al*. 2017). In our previous study (Zill *et al*. 2025), nectar abundance of the dimethylxanthine derivative was associated with female *H. bicruris* host choice. Here, its drought-induced decline in seeds coincided with enhanced larval growth. Although correlative, these findings suggest that methylxanthine metabolism may influence both adult host choice and offspring performance, potentially representing a previously unrecognised chemical link between the mutualistic and antagonistic components of nursery pollination.

### 4.4 Integrating plant sex-specific trait responses across interaction stages: implications for nursery pollination under climatic stress

Overall, our results indicate that water limitation may not necessarily weaken the *S. latifolia–*

*H. bicruris* interaction or shift its net outcome towards antagonism. Instead, climatic stress reorganized resource allocation across different dimensions of the floral phenotype rather than uniformly constraining investment, with consequences for the behavioural decisions through which these phenotypic changes translate into interaction outcomes. On the one hand, water limitation altered multidimensional floral phenotypes and floral sexual dimorphism, particularly nectar chemistry, accompanied by reduced oviposition and a loss of female moths’ oviposition preference for female plants. These responses may reduce seed predation pressure on plants while simultaneously compromising reproductive success of the pollinator, as successful larval development depends on oviposition on seed-producing female plants. On the other hand, changes in seed chemistry under water limitation, particularly reduced defence, coincided with increased larval growth efficiency and earlier pupation at constant seed consumption, potentially benefiting those larvae that successfully develop on water-limited plants without increasing the costs to the plant. Thus, climatic stress may generate opposing effects across interaction stages that cannot be inferred from responses of individual plant traits or from either adult or larval insect responses alone.

By altering the fitness consequences of the interaction for both partners, these responses may also modify reciprocal selective pressures and ultimately their coevolutionary dynamics. Assessing how these effects translate into the net outcome of the interaction under natural conditions will require experiments allowing moths to complete their entire life cycle on plants while accounting for a more complex ecological context, including parasitoids, pathogens, other herbivores, cannibalism, as well as competitive and facilitative interactions between male and female plants.

### 0.8 Acknowledgements

Antonia Wohlmann, Eric Folz, the Botanical Garden Kiel, Maike Pfeiler, Tobias Johannes Demetrowitsch, as well as Katharina Grosser and Verena Zajonc from the Environmental Lab, Kiel University provided technical support. Carolin Böttcher, Saskia Schiffmann, Lisa-Marie Schulz, and Finn Lehmann contributed to data acquisition. Tobias Möckel from the FabLab in Kiel developed a customised camera attachment for light filter switches that was used for the assessment of flower colour composition. This work has been funded by the program for the promotion of young female scientists from the Faculty of Mathematics and Natural Sciences of Kiel University, by institutional overhead funds associated with the German Research Foundation (DFG) projects SCHR 1578/4-1 and SCHR 1578/5-1, and the DAAD Bridge Fellowship Iran. Generative artificial intelligence was used to assist with language polishing and consistency checks of the manuscript.

## 5 Author contributions

Conceptualization: K.S.; Methodology: K.S., T.S., H.B., A.W. and K.Z.; Investigation: R.T., K.Z., A.W., T.S., and H.B.; Validation: K.S., T.S., A.W., H.B., R.T., K.Z., and A.E.; Resources: A.E.; Formal Analysis: K.S.; Data Curation: K.S.; Writing – Original Draft Preparation: K.S.; Writing – Review & Editing: T.S., H.B. and A.E.; Writing – Manuscript Revision: not applicable; Visualization: K.S.; Supervision: K.S.; Project Administration: K.S.; Funding Acquisition: K.S.

