## Supplementary material for "Drought-induced changes in sex-specific floral chemistry reshape interactions with a nursery pollinator across life stages": Figures, Tables, Notes Sx

**Table S1:** See attached xls document.

**Table S2:** Results of statistical analyses testing the effects of watering regime and plant sex on floral traits in Silene latifolia. For univariate response variables, p-values and Wald χ² statistics from type III ANOVAs based on (G)LMMs are shown. For multivariate traits, p-values and pseudo-F statistics from PERMANOVAs are provided. Model interpretation additionally relies on post-hoc comparisons shown in the corresponding figures.

| **(a) Floral traits** | **Watering** | **Plant sex** | **CMI** | **Watering × plant sex** | **Watering × CMI** | **Plant sex × CMI** | **Watering × plant sex × CMI** |
| --- | --- | --- | --- | --- | --- | --- | --- |
| Inflorescence height | **0.000, 13.3** | 0.119, 2.4 | **0.000, 14.3** | 0.422, 0.6 | **0.039, 4.3** | 0.062, 3.5 | 0.484, 0.5 |
| Flower number before hand pollination | **0.000, 14.7** | 0.166, 1.9 | 0.266, 1.2 | 0.735, 0.1 | 0.563, 0.3 | 0.805, 0.1 | 0.859, 0.0 |
| Flower number after hand pollination | **0.001, 11.6** | **0.000, 37.1** | 0.958, 0.0 | 0.110, 2.5 | 0.998, 0.0 | 0.387, 0.7 | 0.667, 0.2 |
| Flower area | **0.000, 14.8** | **0.033, 4.6** | 0.359, 0.8 | 0.571, 0.3 | 0.188, 1.7 | 0.534, 0.4 | 0.749, 0.1 |
| Green:blue reflectance ratio | 0.827, 0.0 | **0.001, 11.6** | 0.224, 1.5 | 0.723, 0.1 | 0.928, 0.0 | 0.665, 0.2 | 0.738, 0.1 |
| Scent metabolome composition | **0.028, 2.4** | **0.000, 6.6** | –^†^ | 0.468, 0.8 | –^†^ | –^†^ | –^†^ |
| Nectar volume | 0.801, 0.1 | **0.000, 53.7** | 0.620, 0.2 | 0.825, 0.0 | 0.894, 0.0 | 0.756, 0.1 | 0.366, 0.8 |
| Nectar metabolome composition | **0.006, 2.9** | **0.001, 3.9** | –^†^ | **0.003, 3.5** | –^†^ | –^†^ | –^†^ |

**Bold lettering** indicates significant effects.

^†^ CMI was not included in the PERMANOVAs because it was measured at the population level and therefore does not vary within the population strata used for permutation.

**Table S3:** Results of statistical analyses testing the effects of watering regime and plant sex on the choice behaviour of Hadena bicruris moths. For univariate response variables, p-values and Wald χ² statistics from type III ANOVAs based on (G)LMMs are shown. For multivariate traits, p-values and pseudo-F statistics from PERMANOVAs are provided. Model interpretation additionally relies on post-hoc comparisons shown in the corresponding figures.

| **(b) Pollinator choice behaviour** | **Preference (intercept)** | **Treatment (watering or plant sex)** | **Moth sex** | **Plant population** | **Treatment × moth sex** | **Treatment × plant population** | **Moth sex × plant population** | **Treatment × moth sex × plant population** |
| --- | --- | --- | --- | --- | --- | --- | --- | --- |
| Feeding (W vs. L) | 0.256, 1.28 | 0.560, 0.3 | 0.228, 1.4 | 0.905, 0.0 | 0.209, 1.6 | 0.325, 1.0 | **0.040, 4.2** | 0.295, 1.1 |
| Oviposition (W vs. L) | **0.034, 4.50** | 0.589, 0.3 | –^†^ | 0.256, 1.3 | –^††^ | 0.691, 0.2 | –^†^ | –^†^ |
| Feeding (F vs. M) | 0.252, 1.31 | 0.716, 0.1 | 0.256, 1.3 | **0.016, 5.8** | 0.398, 0.7 | 0.334, 0.9 | 0.44, 0.6 | 0.679, 0.2 |
| Oviposition (F vs. M) | **0.001, 10.37** | **0.02, 5.5** | –^†^ | 0.348, 0.9 | –^†^ | 0.467, 0.5 | –^†^ | –^†^ |

**Bold lettering** indicates significant effects.

^†^ Moth sex not included as only female moths can oviposit.

**Table S4:** Results of statistical analyses testing the effects of watering regime and further covariates on seed traits in Silene latifolia and Hadena bicruris larval performance and consumption rate. For univariate response variables, p-values and Wald χ² statistics from type III ANOVAs based on (G)LMMs are shown. For multivariate traits, p-values and pseudo-F statistics from PERMANOVAs are provided.

|  | **Water** | **CMI** | **Water × CMI** | **Initial age larva** | **Consumed seed mass** | **Water × Consumed seed mass** |
| --- | --- | --- | --- | --- | --- | --- |
| Seed set (conditional) | 0.069, 3.3 | 0.570, 0.3 | 0.312, 1.0 | – | – | – |
| Seed set (zero inflation) | **0.048, 3.9** | –^†^ | –^†^ | – | – | – |
| Seed mass per area | **0.036, 4.4** | 0.556, 0.3 | 0.881, 0.0 | – | – | – |
| Seed germination rate | 0.854, 0.0 | 0.295, 1.1 | 0.486, 0.5 | – | – | – |
| Mean seed germination time | 0.655, 0.2 | 0.467, 0.5 | 0.093, 2.8 | – | – | – |
| Seed metabolome composition | 0.174, 1.32 | –^††^ | –^††^ | – | – | – |
| Relative growth rate larva | **0.018, 5.6** | 0.319, 1.0 | 0.859, 0.0 | 0.984, 0.0 | 0.056, 3.6 | **0.000, 21.3** |
| Time to pupation | **0.049, 3.8** | 0.913, 0.0 | 0.072, 3.2 | **0.005, 7.7** | – | – |
| Pupation rate | 0.340, 0.9 | 0.205, 1.6 | 0.431, 0.6 | 0.781, 0.1 | – | – |
| Total seed mass consumed | 0.204, 1.6 | 0.653, 0.2 | 0.656, 0.2 | 0.594, 0.3 | – | – |

**Bold lettering** indicates significant effects.

^†^ Inclusion of fixed effect did not improve model AIC and was thus neglected for parsimony.

^††^CMI was not included in the PERMANOVA because it was measured at the population level and therefore does not vary within the population strata used for permutation.

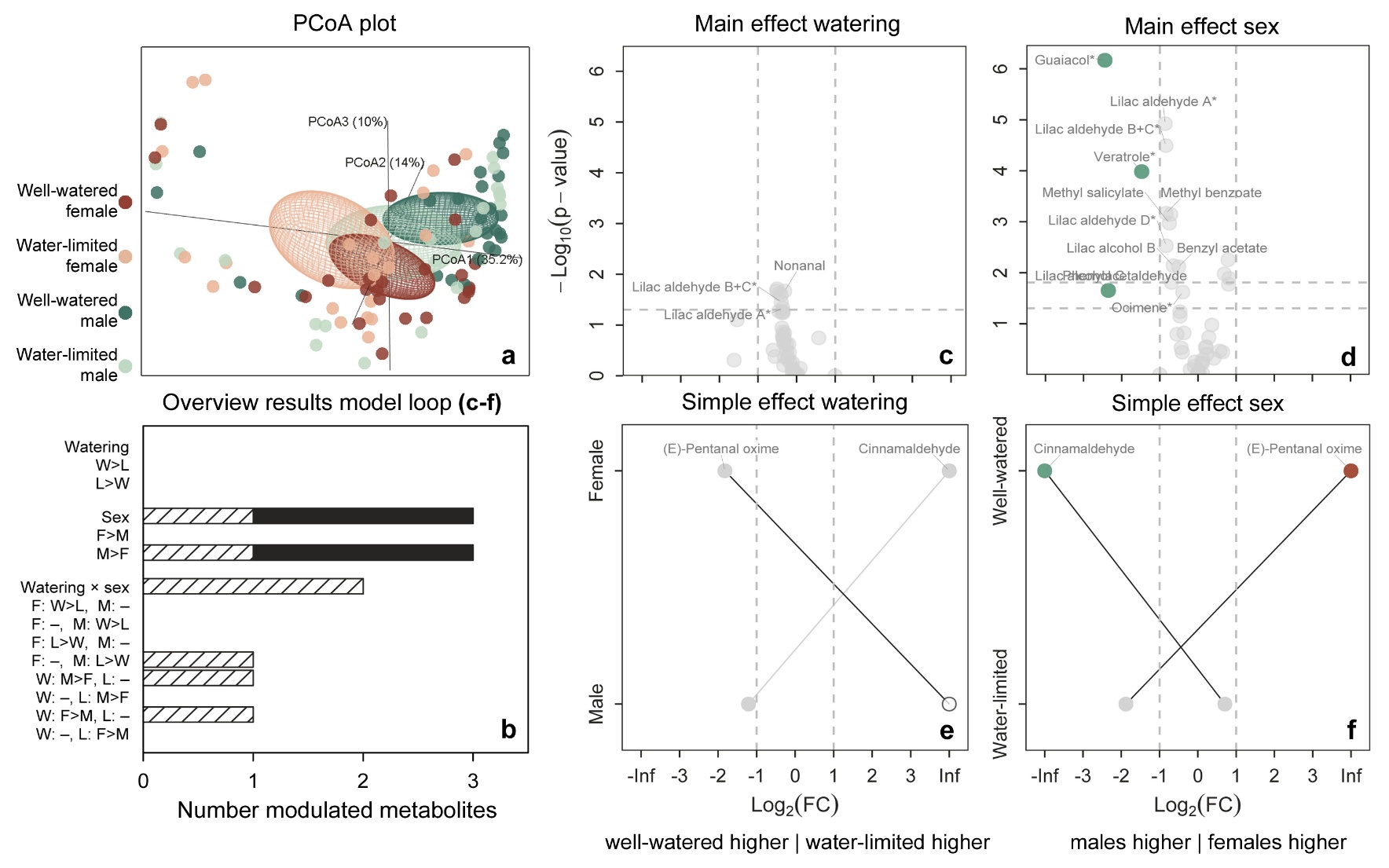

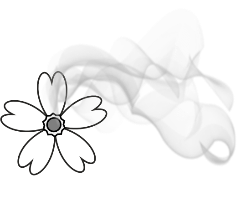
**Figure S1:** Effects of watering regime and plant sex on floral scent composition in Silene latifolia. **(a)** 3D PCoA plot with points representing well-watered females (W–F, dark red), water-limited females (L–F, light red), well-watered males (W–M, dark turquoise), and water-limited males (L–M, light turquoise). Ellipsoids represent 95% confidence regions around group centroids. **(b)** Bar plot summarizing model-loop results as the number of features modulated by each effect and direction (p_RAW_ / p_FDR_ <0.05 [dashed / filled] and |log_2_FC| > 1). **(c***–***d)** Volcano plots showing the main effects of watering regime and plant sex for features not additionally involved in a significant interaction. Modulated features (p_RAW_ / p_FDR_ <0.05 [lower / upper horizontal dashed line] and log_2_FC < -1 / > 1 [left / right vertical dashed line]) are coloured: higher in well-watered plants= dark grey, higher in water-limited plants = white, higher in females = red, higher in males = turquoise. Interaction effect volcano plots show **(e)** the simple effect of watering regime within females and males, and **(f)** the simple effect of plant sex within well-watered and water-limited plants. Identical features within these group comparisons are connected by lines. For further details on the identity of metabolic features, see Table S1a. N = 5 populations × 2 sexes × 2 watering regimes × 4–5 replicates = 95.

Note S1: Extended Methods

Plant and pollinator cultivation (Chapter 2.2)

*S. latifolia* plants from the F1 generation were raised in 360 mL pots (TKS2 Instant Plus Substrate, Floragrad, Oldenburg, DE; supplemented with 4.5 g L⁻¹ Basacote® Plus 6M, Compo Expert GmbH, Münster, GE) at 25 °C / 15 °C under a 16 h light / 8 h dark cycle in climate chambers (HGC 1514, Weiss Technik, Resikirchen, DE; equipped with LED C90 NS12, Valoya, Helsinki, FI). After flowering onset, we selected 10 female and 10 male plants per population for the experiment and transferred them to 3 L pots. Additional plants were kept to maintain our laboratory *H. bicruris* population. Plants were watered to keep soil moisture at optimal levels and treated prophylactically with biological pest control (Katz Biotech GmbH, Baruth, DE) during an initial eight-week growing period.

Adult *H. bicruris* moths were collected from several wild *S. latifolia* populations in the Kiel region, Germany (54°19′ N, 10°8′ E) in 2022 and maintained as a mixed breeding stock in ventilated transparent boxes (20 × 30 cm) furnished with egg cartons and tissue paper. Boxes were kept in a climate chamber (MLR-352H, PHC Corporation, Ōizumi-machi, JP) at 25 °C / 15 °C, 60 % relative humidity, and a 16 h light / 8 h dark cycle. Adults were fed a honey-water solution (6 g organic flower honey in 14 mL distilled water) and additionally supplied with fresh male and female S. latifolia flowers, as reproduction did not occur without these. Flowers were harvested from a pool of food plants from all studied populations and offered individually in water-filled tubes. Eggs laid throughout the boxes were collected and incubated on moist filter paper in individual containers to prevent cannibalism. After hatching, larvae were individually reared on an artificial diet (Elzinga *et al.*, 2002) supplemented with S. latifolia seeds (20 mL per 500 mL diet, study population mix) to increase acceptance. Pupae were transferred to individual ventilated 50 mL containers containing moist sterile sand, and newly emerged adults were moved back to the breeding boxes and feeding regime described above.

Visual flower traits: Colour composition and area (Chapter 2.4.1)

Photos of flowers were taken under a metal halide bulb (70 W Sunray, ExoTerra, Holm, GE) approximating the spectral composition of natural sunlight. One flower was placed in a black ethylene vinyl acetate platform equipped with two reflectance standards (PTFB 10%; Spectralon 99% Labsphere, Chester-le-Street, UK). Raw images were captured with a digital camera (Samsung NX1000, Suwon, KR) converted to full-spectrum sensitivity (300 to 1000 nm) and fitted with a UV-sensitive lens (Nikon EL 80-mm, Tokyo, JP). We acquired both visible and UV images by fitting either a UV and infrared blocking filter or a UV-pass plus IR-block filter, respectively (UV/IR Cut filter: transmittance 400 to 700 nm, U-filter: transmittance 300 to 400 nm, Baader Planetarium, Mammendorf, DE). Images were processed using the Multispectral Image Calibration and Analysis (MICA) Toolbox plugin (Troscianko and Stevens, 2015) in ImageJ 1.47 t (Rueden *et al.*, 2017). Petals were selected for the quantification of area, as well as green, blue and UV reflectance. As *S. latifolia* petals absorb more than 95% of the UV light at the display side (Schrieber *et al.*, 2021; Zill *et al.*, 2025), and as nocturnal moth vision is often insensitive to red wavelengths (Kelber *et al.*, 2003), we focused on blue and green reflectance only.

**Floral scent metabolome (chapter 2.4.2)**

Samples were analysed in one run in fully randomised order by thermal desorption gas chromatography mass spectrometry (TD-GC-MS: TD-30R, GCMS-2020NX, Shimadzu, Kyoto, JP). Trapped VOCs were desorbed from PDMS tubes for 8 min at 230 °C under a helium flow of 60 mL min**^−^**¹ and adsorbed on a Tenax TA filled cryo-trap at **−**20 °C. From there, compounds were desorbed at 250 °C for 3 min, injected into the GC with split ratio of 3:3, and separated under a helium flow of 1.6 mL min**^−^**¹ on a RTX-5MS column (30 m × 0.25 mmx 0.25µm , Restek, Bad Homburg, DE). The oven programme was: 40 °C for 5 min, 40 °C to 125 °C at 10 °C min**^−^**¹, 5 min hold, 10 °C to 280 °C at 30 °C min**^−^**¹, 1.33 min hold. Mass spectra from 30 to 400 m/z were acquired in scan mode with an event time of 0.10 sec. An alkane mix (C_8_ to C_20_, Sigma-Aldrich, Darmstadt, DE) was run under identical conditions to calculate linear retention indices (RI) for the targeted compounds. Compounds were identified by matching RI and mass spectra to synthetic standards, published references (Dötterl *et al.*, 2005; Mamadalieva *et al.*, 2014), and library entries (NIST17: Smith *et al.* 2004, Pherobase: El-Sayed 2011). Controls (16 units without flowers) and blanks (solvent- and heat-cleaned PDMS tubes) were used to identify and exclude contaminants. All compounds were semi-quantified using the total ion chromatogram peak intensities.

**Floral nectar metabolome (chapter 2.4.3)**

2 µL of extract were injected at 30 °C into a UHPLC (Nexara x3, Shimadzu, Kyoto, JP) containing a reversed phase column (100 × 2.1 mm, 1.7 µm, Luna Omega Polar C_18_, Phenomenex, Aschaffenburg, DE). The solvent system was composed of (A) water with 0.1% formic acid, and (B) acetonitrile with 0.1 % formic acid (all ROTISOLV Ultra LC-MS grade, Carl Roth, Karlsruhe, DE). The following gradient program was run: 0.3 mL min**^−^**¹ flow, starting at 5 % B, ramped to 95 % B over 20 min, held 5 min, then re-equilibrated at 5 % B for 6 min. Chromatographically separated analytes entered a qTOF-MS/MS (x500R, Sciex, Darmstadt, DE) with the following parameters: ion source gas 45 psi (N₂), curtain gas 35 psi, CAD gas 7 psi, source temperature 500 °C, ion spray voltage 5500 V (positive mode) or **−**4500 V (negative mode), MS1 scan range 100-1000 m/z, data-dependent MS2 trigger with 40 eV collision energy and a scan range of 50-1000 m/z. Data were converted to .mzML format and processed in mzMine (Schmid *et al.*, 2023). Features with a maximum m/z difference of 0.001 Da and retention time difference of 0.1 min were considered identical. For features detected in both the hydrophilic and lipophilic fractions, signal intensities were summed across fractions.

For nectar feature annotation, the acquired MS/MS spectra were compared against public mass spectral libraries (GNPS: Wang *et al.* 2016, MassBank: MassBank consortium and its contributors 2024) using the NIST MS search algorithm (cosine similarity of 0.6) and the MassBank formula (cosine similarity of 0.5). Discriminatory features that remained unknown were further annotated using COSMIC and CSI:FingerID according to the SIRIUS software package (Hoffmann *et al.*, 2022). Following the first comprehensive description of nectar chemistry in the ecological model system *S. latifolia* based on a single population (Zill *et al.*, 2025), we now report the first cross-population assessment.

**Statistical analyses of floral VOCs, nectar and seed metabolomes for discriminatory feature selection (chapter 2.6.2)**

To determine which individual features contributed to treatment- and sex-related variation, we applied a feature-level modelling approach. This approach is comparable to a volcano plot and simultaneously allows the statistical assessment of interaction effects among plant sex and watering treatment, as well as accounting for the highly heterogeneous distributional properties of GC/LC–MS metabolite intensities. Instead of applying two-group significance tests, we entered all metabolites as responses into a (G)LMM. This procedure followed a two-step logic: (1) identification of the best-fitting model and (2) downstream analysis of that model. Model selection was based on residual diagnostics and likelihood criteria, accommodating heterogeneity in metabolite intensities, including distributional shape, skewness, heteroscedasticity, and a high proportion of values below the detection limit. For each metabolite, we fitted a set of candidate (G)LMMs in a custom R loop (R-package “glmmTMB”, Brooks *et al.*, 2017), all with watering regime, plant sex, and their interaction as fixed effects and population as random effect. CMI was omitted as fixed effect for parsimony, as it explained only little variation in overall metabolome composition ***(Supplementary Note S1)***. Candidates included Gaussian models on the raw response scale and on transformed response scales (log1p, square root), Gamma models with log link, and Bernoulli models (above vs. below the detection limit). For Gaussian and Gamma models, we compared constant vs. group specific dispersion (by watering regime and plant sex). The candidates were evaluated using DHARMa residual diagnostics (R-package "DHARMa", Hartig 2022). Gaussian and Gamma models were assessed with tests for residual uniformity, dispersion, and homogeneity of residuals across groups, whereas Bernoulli models were assessed only for residual uniformity. Selection proceeded by diagnostic filtering, preferring models with fewer violations, followed by AIC among diagnostically equivalent models of comparable response scales and error distributions. Continuous models were preferred whenever diagnostically acceptable. Bernoulli models were considered only when all continuous candidate models failed residual diagnostics. Statistical inference was performed only for the best supported model per metabolite. Downstream analysis of the selected best models was performed with the R-package “emmeans” (Lenth 2024). Estimated marginal means were computed and regridded to the response scale, such that all subsequent contrasts and statistical inference were performed in the original response units. Contrasts representing the overall effects of sex, watering treatment, and their interaction were calculated from these response-scale estimates. When interactions were significant, predefined contrasts between treatment combinations were additionally evaluated. P values were adjusted for multiple testing separately for main effects and interactions using the Benjamini–Hochberg procedure. Compounds with large magnitude changes (|log_2_FC| > 1) and p_FDR_<0.05 were prioritized in the interpretation of data. While FDR correction and fold-change thresholds are useful statistical tools for identifying potentially important features among a multitude of unknown metabolites, biologically relevant effects may also arise from small changes in compound abundance. We therefore additionally report and discuss nominally significant and low-magnitude changes for identified compounds with known ecological functions

Note S2: Effects of climate moisture index on plant–insect interaction responses to watering regime

In contrast to watering regime and plant sex, CMI had rather limited effects on interaction-associated traits in *S. latifolia* and *H. bicruris* (Manuscript Table 1–2). Inflorescence height was significantly shaped by the interaction watering regime × CMI, with an increasingly negative effect of water limitation from the drier towards the wetter end of the CMI gradient (Table S2, Note S2 Figure 1a). Inflorescence height was also shaped by the interaction between plant sex and CMI, with females having taller inflorescences than males at the drier end of the CMI gradient, while the direction of this sexual dimorphism reversed towards the wetter end (Table S2, Note S2 Figure 1b). Although CMI was included in all univariate trait models, the random effect of population still accounted for substantial variance in several traits, indicating pronounced population divergence beyond the climatic gradient. For floral and seed chemistry, minor effects of population-level CMI were detected only for floral scent, with generally lower abundances of several VOCs towards the wetter end of the CMI gradient (Note S2 Table 1–2).

*Note S2 Figure 1***:** Interactive effects of watering regime plant sex and summer climate moisture index on inflorescence height in Silene latifolia. Points represent observations for female (red) and male (turquoise) plants under well-watered (dark) and water-limited (L–bright) conditions. The corresponding lines represent predictions from the (G)LMM with 95% confidence intervals. The results of omnibus tests (type III Anova based on Wald χ²-tests) are denoted at the top of each plot while results of post – hoc of partial slopes are denoted within plots.

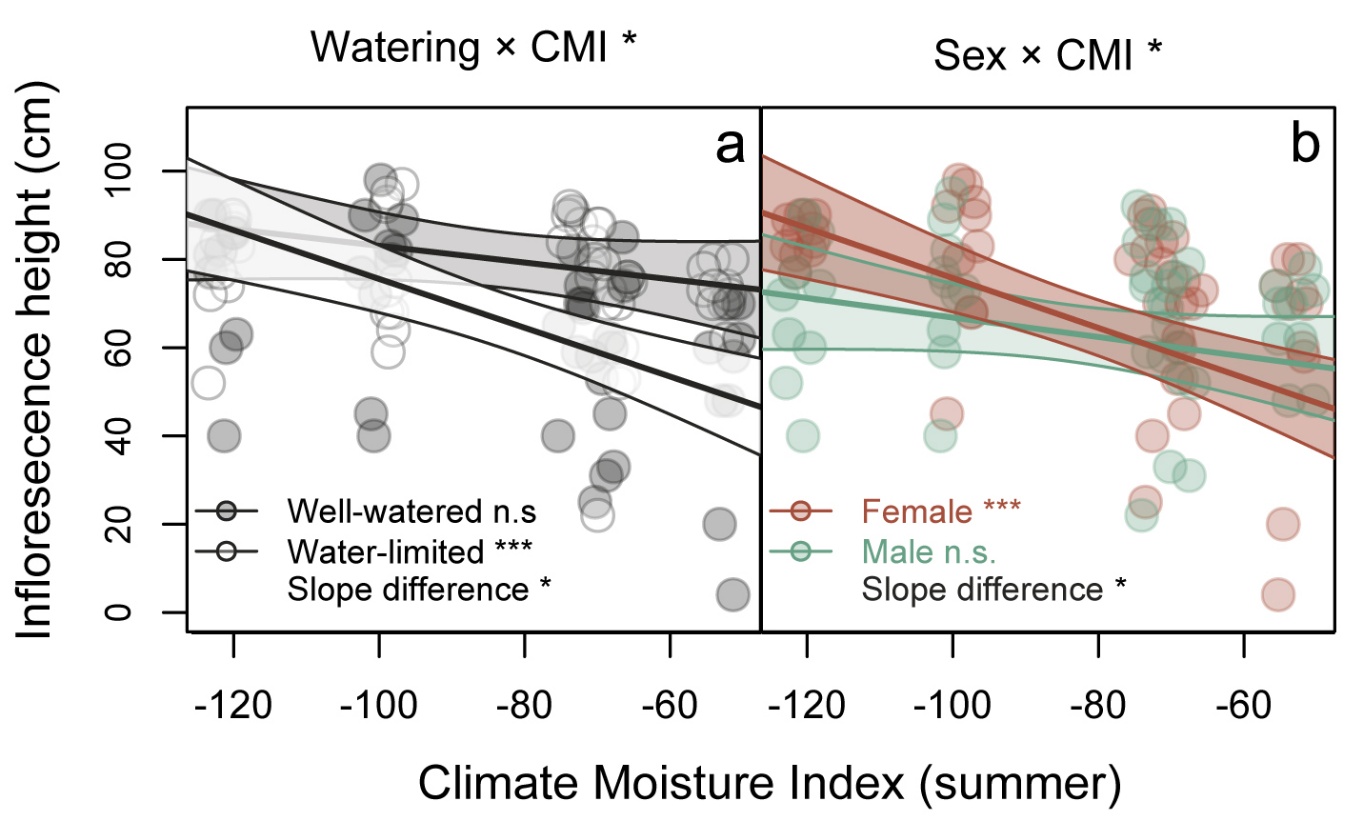

*Note S2* **Table 1:** Overview of PERMANOVA results for the individual and combined effects of watering regime, plant sex and CMI on the composition of the floral scent, nectar and seed metabolome. Note that metabolome data were averaged on the population × watering regime × plant sex level, as PERMANOVA cannot model covariates (i.e., CMI) that are constant within a stratum (population). As the effects of CMI were at most minor, it was neglected for parsimony and interpretability of all metabolome datasets in the full article. However, *Note S2 Table* **2** provides the results of floral scent metabolome analyses incorporating CMI as a main effect as indicated by the PERMANOVA model.

| **Matrix** | **Output value** | **Watering regime** | **Plant sex** | **CMI** | **Watering regime × plant sex** | **Watering regime × CMI** | **Plant sex × CMI** | **Watering regime × plant sex × CMI** |
| --- | --- | --- | --- | --- | --- | --- | --- | --- |
| Floral VOC | R² | 0.059 | **0.193** | **0.119** | 0.023 | 0.066 | 0.071 | 0.023 |
|  | pseudo F | 1.584 | **5.199** | **3.217** | 0.627 | 1.768 | 1.922 | 0.606 |
|  | p | 0.166 | **0.002** | **0.017** | 0.693 | 0.122 | 0.102 | 0.701 |
| Floral nectar | R² | 0.104 | 0.086 | 0.087 | **0.143** | 0.014 | 0.017 | 0.040 |
|  | pseudo F | 2.454 | 2.027 | 2.055 | **3.382** | 0.334 | 0.396 | 0.937 |
|  | p | 0.066 | 0.113 | 0.105 | **0.026** | 0.876 | 0.830 | 0.430 |
| Seed | R² | 0.087 | not tested | 0.135 | not tested | 0.067 | not tested | not tested |
|  | pseudo F | 0.735 | not tested | 1.144 | not tested | 0.570 | not tested | not tested |
|  | p | 0.798 | not tested | 0.315 | not tested | 0.911 | not tested | not tested |

*Note S2* **Table 2:** Overview of CMI effects on the composition of floral scent. In accordance with the results of PERMANOVA (Table S2a) CMI was included as main effect into the floral scent metabolite model loop in an additional analysis. Only six of 45 floral VOC correlated significantly with CMI.

| **Compound** | **p** | **pFDR** | **Effect direction** |
| --- | --- | --- | --- |
| 1-Dodecanol | 0.044 | 0.353 | Reduced toward wetter end of CMI gradient |
| Benzenepropanol | 0.006 | 0.097 | Reduced toward wetter end of CMI gradient |
| Ethyl benzoate | 0.030 | 0.339 | Reduced toward wetter end of CMI gradient |
| Phenylacetaldehyde† | 0.001 | 0.028 | Increased toward wetter end of CMI gradient |
| Unknown 1 | 0.047 | 0.353 | Reduced toward wetter end of CMI gradient |
| Veratrole† | 0.000 | 0.011 | Reduced toward wetter end of CMI gradient |

†Compound is known to elicit antennal and behavioural responses in Hadena bicruris according to Dötterl et al. (2006)

Note S3: Integrated evidence for methylxanthine metabolism in *Silene latifolia*

Summary

The novel structurally unresolved dimethylxanthine derivative previously discovered in *S. latifolia* nectar (Zill *et al.*, 2025), was again detected in nectar and additionally in seeds in the present study Moreover, xanthosine, a precursor of methylxanthines, occurred at high relative abundance in seeds. (chapters 3.1***–***2, Figures 3 and 6, Table S1b***–***c). To further examine this recurring pattern, we analysed hydrophilic extracts from mature seeds and germinated seedlings with green leaves by D1 ^1^H-NMR, which independently confirmed the occurrence of xanthosine (12.7 nmol mg^-1^) and a dimethylxanthine compound (6.1 nmol mg^-1^). To further evaluate the occurrence of methylxanthine metabolism in *S. latifolia*, we annotated the predicted proteome of the recently published reference genome (Moraga *et al.* 2026) using KOfamScan. Multiple proteins were assigned to KEGG orthologues of caffeine synthases (S-adenosyl-L-methionine-dependent methyltransferases), providing genomic evidence for the enzymatic machinery required for N-methylation of xanthine derivatives. These predictions are consistent with the structure of the novel dimethylxanthine derivative detected in *S. latifolia* and provide independent genomic support for the presence of methylxanthine metabolism. A detailed summary of these complementary lines of evidence is provided below.

**Evidence from UHPLC-qTof-MS/MS analyses and chemoinformatics**

Metabolomic analysis of *S. latifolia* nectar and seeds using UHPLC-qToF-MS/MS detected a shared metabolic feature with an *m/z* of 351.0733 and a retention time of 1.06 min. Its MS^2^ spectrum did not yield a match to compounds in publicly available spectral databases (***Note S3 Figure 1***). Further analysis using MZmine and *in silico* annotation tools implemented in the SIRIUS platform predicted the molecular formula C₁₄H₁₅N₄O₅S. Combining the predicted molecular formula with the observed fragmentation pattern enabled a tentative *de novo* annotation of the feature as a substituted methylxanthine derivative. However, the MS^2^ spectrum contained no sufficiently diagnostic fragments beyond those attributable to the xanthine-related core structure to resolve the identity of the substituent. Based on the predicted molecular formula, the substituent at the N9 position may contain a cyclic moiety bearing additional hydroxy and thiol functionalities (***Note S3 Figure 2***).

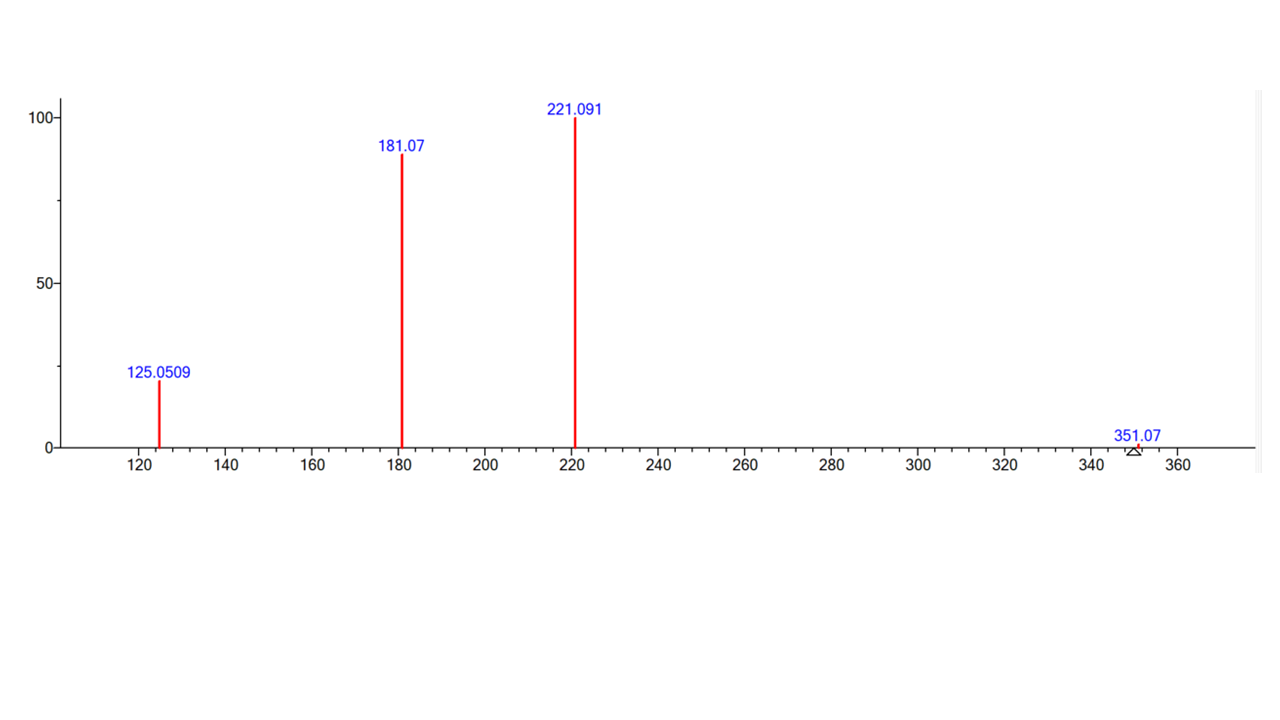

**Note S3 Figure 1:** *MS^2^ spectrum of the putative methylxanthine derivative acquired in positive ionization mode at a collision energy of 40 eV. Precursor ion: m/z 351.0733; RT: 1.06 min.*

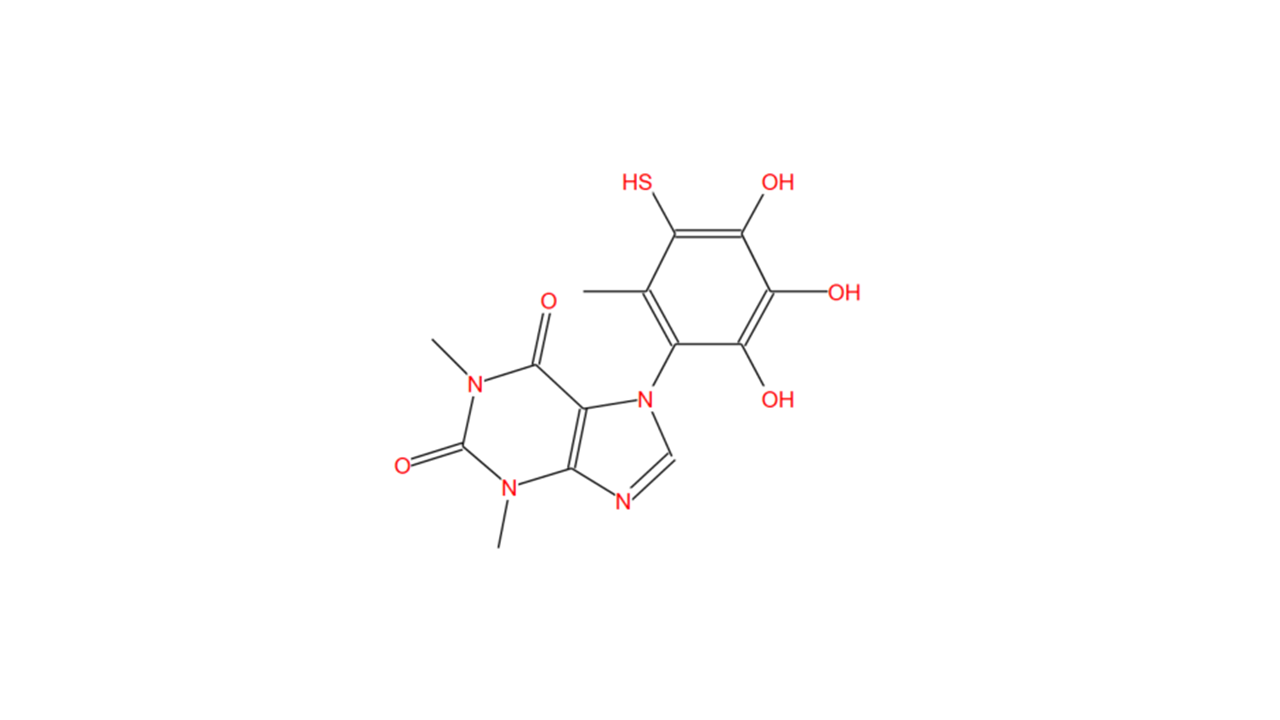

**Note S3 Figure 2:** Proposed de novo structure of the methylxanthine derivative. The structure of the substituent at the N9 position is hypothetical and is based primarily on the predicted molecular formula. The available MS^2^ data do not provide sufficient evidence to establish its connectivity or exact structure.

**Evidence from 1D ^1^H -NMR**

The abundance of the dimethylxanthine derivative was further evaluated using 1D ^1^H NMR. Seeds and germinated seedlings with green leaves from all study populations were used because they provided sufficient biomass, unlike nectar. Samples were frozen in liquid nitrogen, ground to powder, and 100 mg seed or 150 mg seedling powder was extracted in 1 mL acetonitrile/ddH₂O (1:1) for 2 h at 12 °C. After centrifugation (14,000 × *g*, 10 min), supernatants were snap-frozen, lyophilised, and redissolved in 600 μL NMR buffer (90 mM sodium phosphate, 10% D₂O, 4.6 mM DSS, pH 7.4). Water-suppressed 1D ^1^H NMR spectra were recorded with 128 scans on a Bruker Avance 600 MHz spectrometer equipped with a TCI cryoprobe using the zgesgp pulse sequence. Spectra were processed in TopSpin and analysed and quantified using Chenomx NMR Suite v10.1.

The seed metabolome contained xanthosine, identified by assignment of the purine C8 and ribose C1 and C3 proton shifts against the Chenomx reference spectrum (***Note S3 Figure 3–4***). The predicted C4 and C5 ribose signals overlapped with resonances of other metabolites, preventing unambiguous assignment, while the predicted C2 signal (4.38 ppm) was obscured by water suppression around the water resonance (4.7 ppm). Although xanthosine can occur as a degradation product, its high abundance of 12.7 ± 2.6 nmol mg⁻¹ in seeds suggests that it may instead serve as a precursor for methylxanthine biosynthesis. The methylxanthine derivative, previously detected in seeds by UHPLC-qToF-MS/MS, was not detectable by 1H NMR, indicating that its abundance was below the detection limit of solution NMR (~1 μM). The germling metabolome on the other hand lacked characteristic resonances of xanthosine, but C8-, N1(CH_3_)- and N3(CH_3_)-proton signals corresponding to a methylxanthine metabolite were identified (***Note S3 Figure 5–6***). The chemical shifts were assigned based on the reference spectrum of trimethylxanthine (caffeine, C8-proton: 7.89 ppm, N1(CH_3_)-proton: 3.52 ppm, N3(CH_3_)-proton: 3.34 ppm). The metabolite was present at 6.1 ± 1.1 nmol mg⁻¹ fresh seedling material.

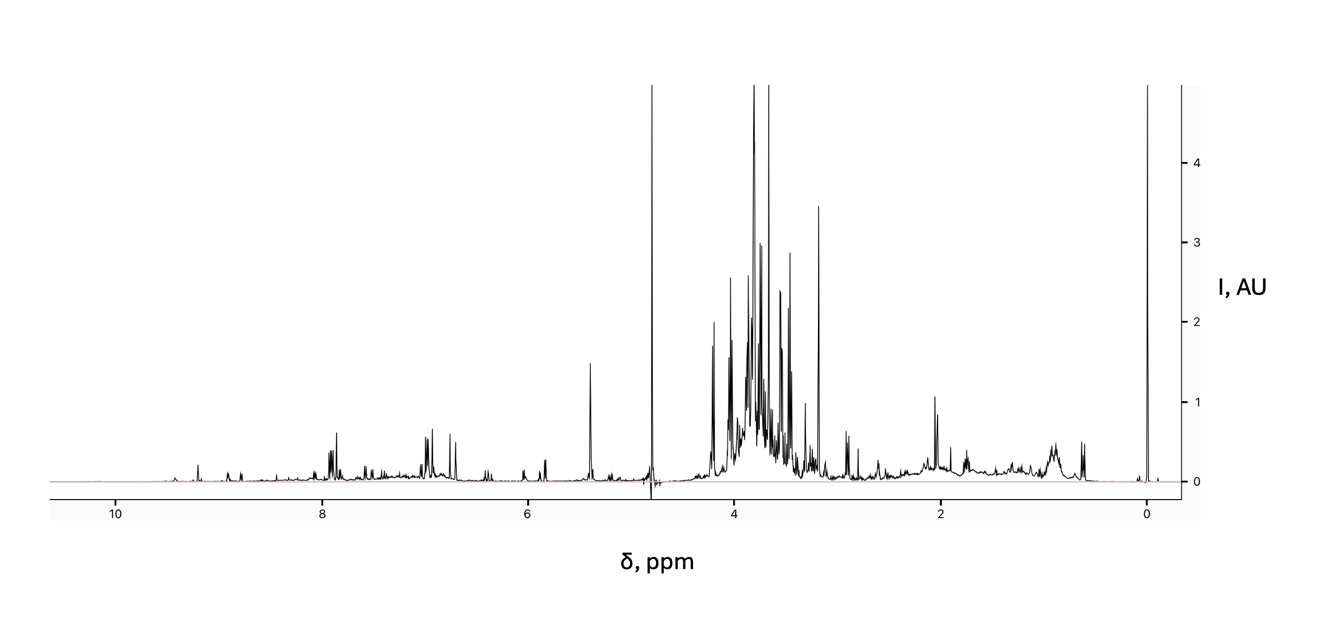

**Note S3 Figure 3:** Representative 1D-1H NMR spectrum of S. latifolia seed hydrophilic metabolome extract.

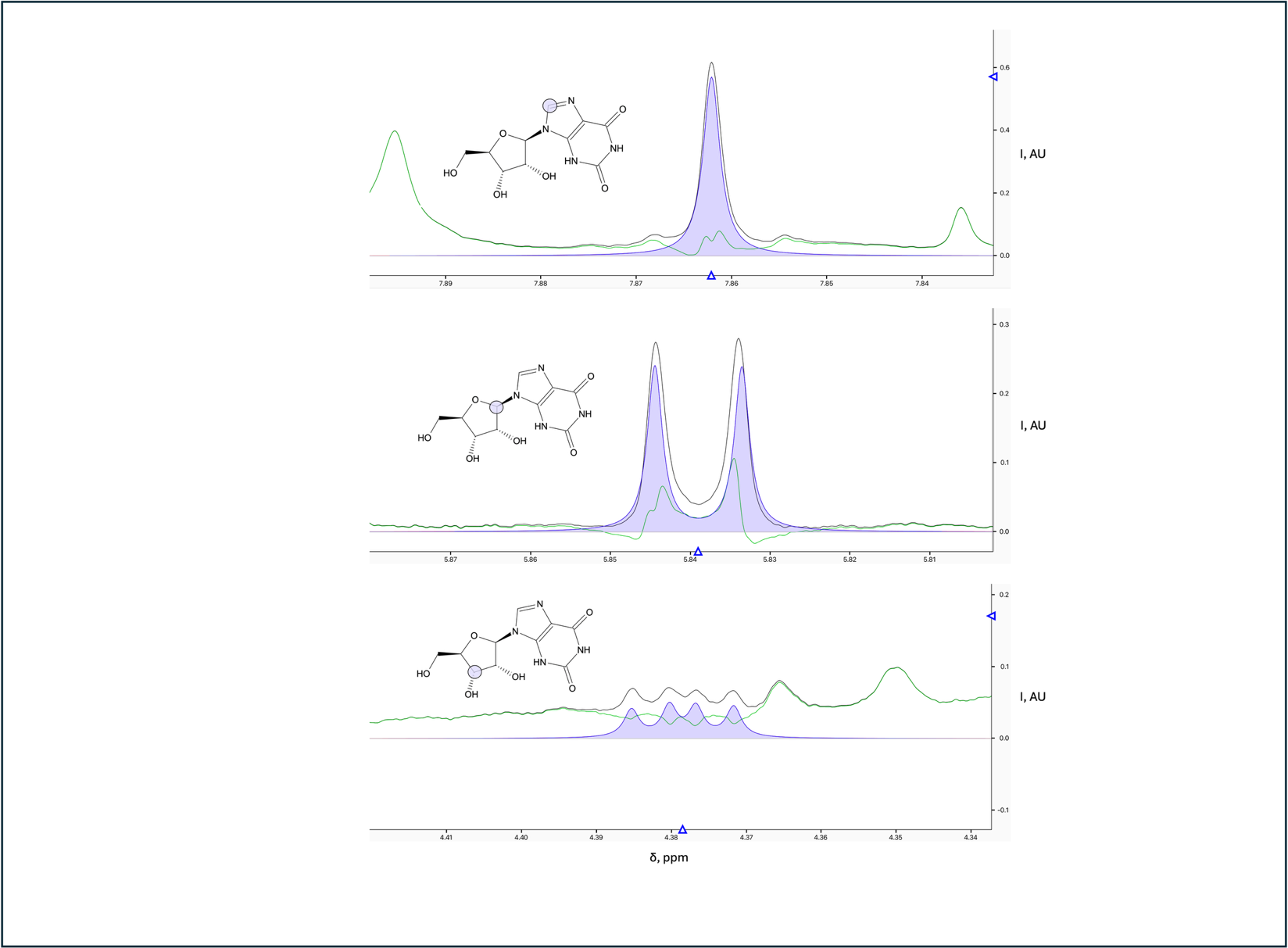

**Note S3 Figure 4:** Spectral peaks of the S. latifolia hydrophilic seed metabolome extract that could be unambiguously assigned to Xanthosine protons. Peaks of the reference spectrum of Xanthosine (blue) were fitted to the Lorentzian lineshapes of the spectrum (black). A subtraction line (green) reports the predicted spectral shape subtracting the signals coming from the fitted reference compound.

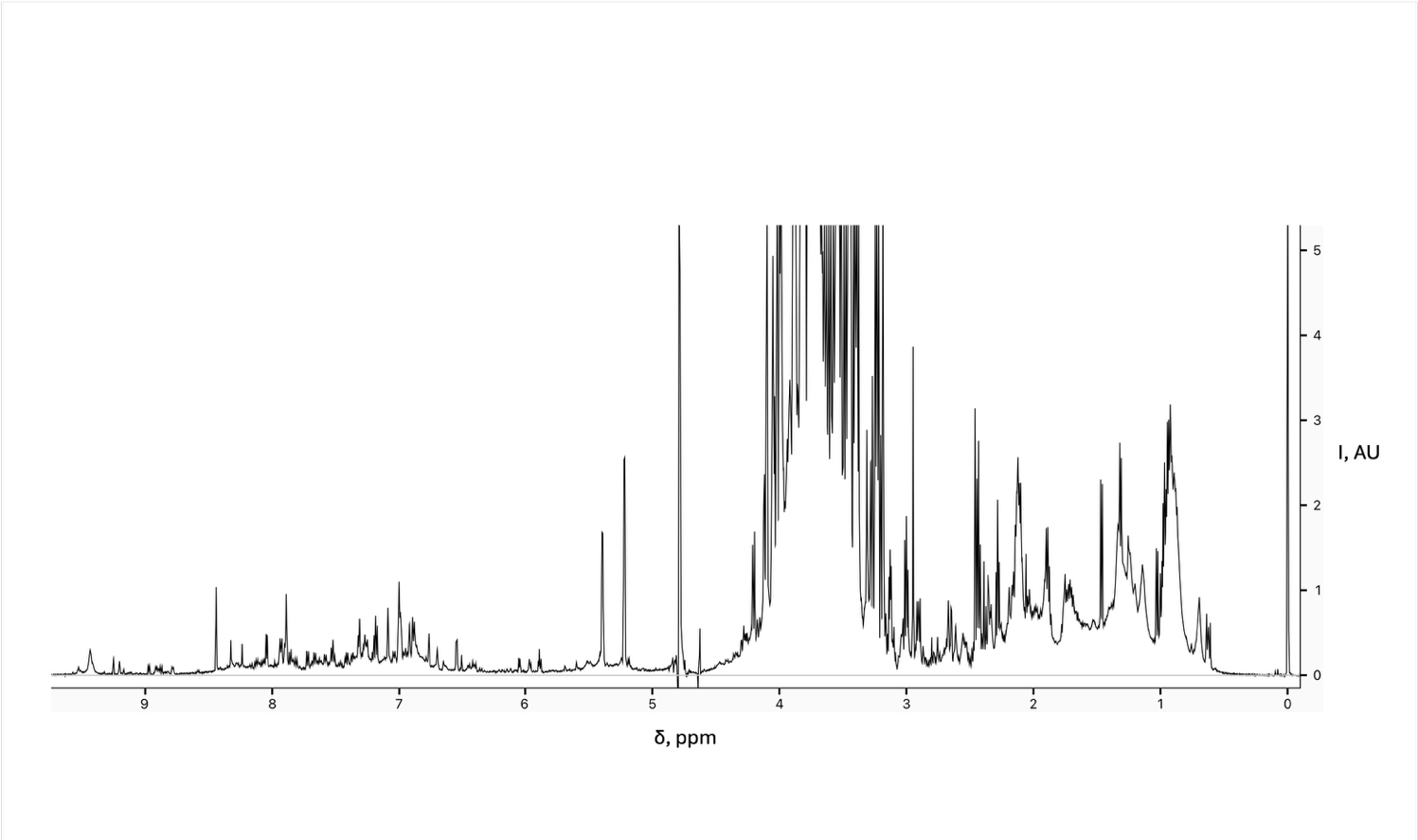

**Note S3 Figure 5:** Representative 1D-1H NMR spectrum of S. latifolia germling hydrophilic metabolome extract.

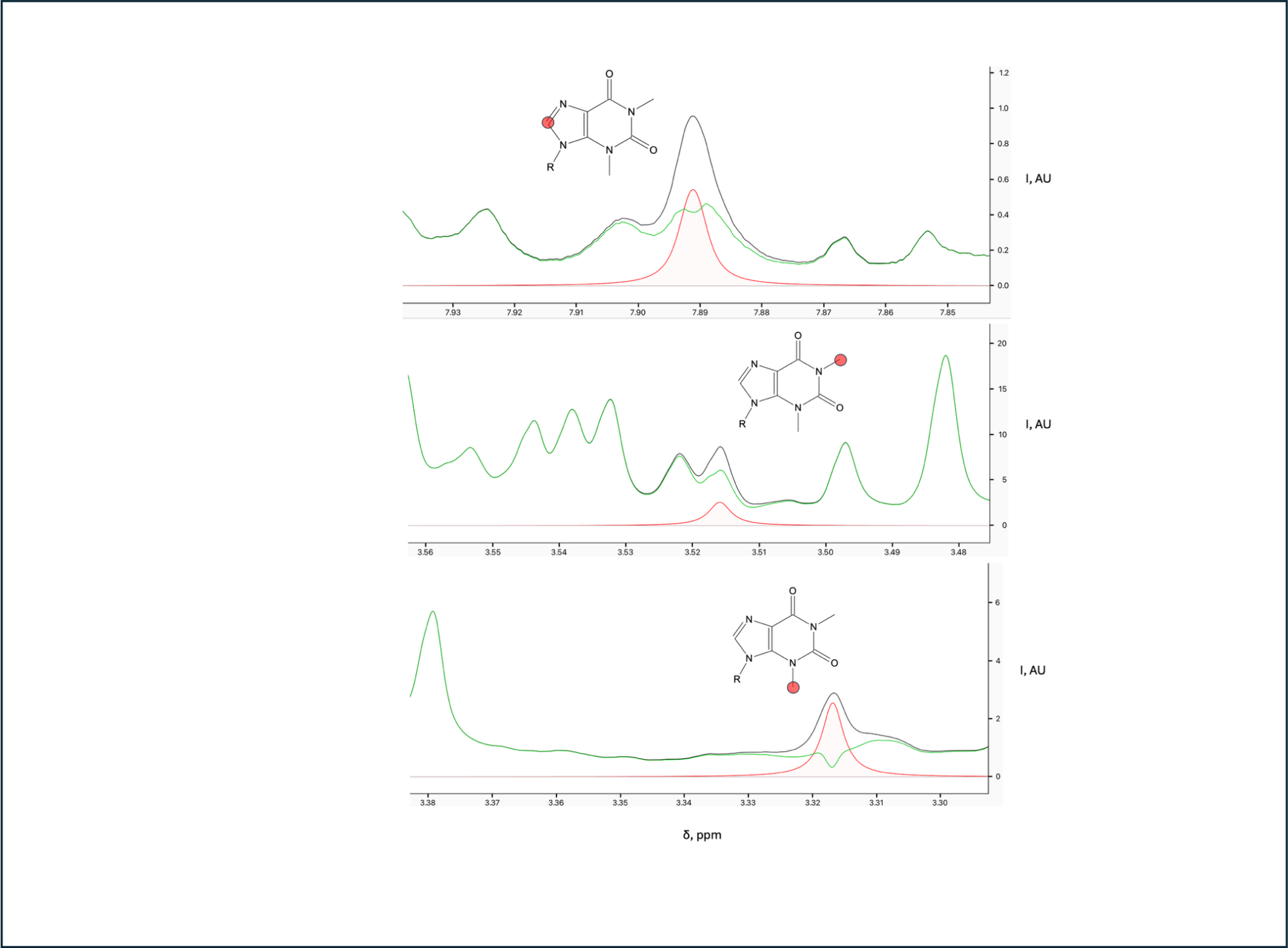

**Note S3 Figure 6:** Spectral peaks of the S. latifolia hydrophilic metabolome extract that could be assigned to methylxanthine proton resonances, based on the reference spectrum of caffeine. Assigned peaks (red) were fitted to the Lorentzian lineshapes of the spectrum (black). A subtraction line (green) reports the predicted spectral shape subtracting the signals coming from the fitted reference compound.

**Evidence from comparative genome annotation**

A recent study integrating short-read, long-read and high-C genomic DNA sequencing allowed for the chromosome scaffolded assembly of the *Silene latifolia* genome (Moraga *et al.* 2026). The assembled genome was annotated by NCBI Eukaryotic Genome Annotation Pipeline in multiple steps, including alignments of RNA-seq data from multiple studies for gene prediction. The published list of predicted proteins was downloaded and annotated for KEGG Orthologs using KofamScan v1.3.0 with the eukaryote profile and mapper function (Ogata *et al.*, 1999; Aramaki *et al.*, 2020). The list of KO-assigned genes was visualized using the KEGG Mapper Reconstruct tool and inspected for modules of methylxanthine metabolism (***Note S3 Figure 7***). We identified three KEGG orthologs for N-methylxanthine methylating S-adenosyl-L-methionine dependent methyltransferases (KO K12731), often referred to as Caffeine Synthases, which can methylate the N1 and N3 positions of methylxanthines (Uefuji *et al.*, 2003). As the methylxanthine compound we observed in our experimental metabolomics analyses is methylated at both N1 and N3, the presence of these methyltransferases is an independent layer of evidence that *S. latifolia* can produce and modify
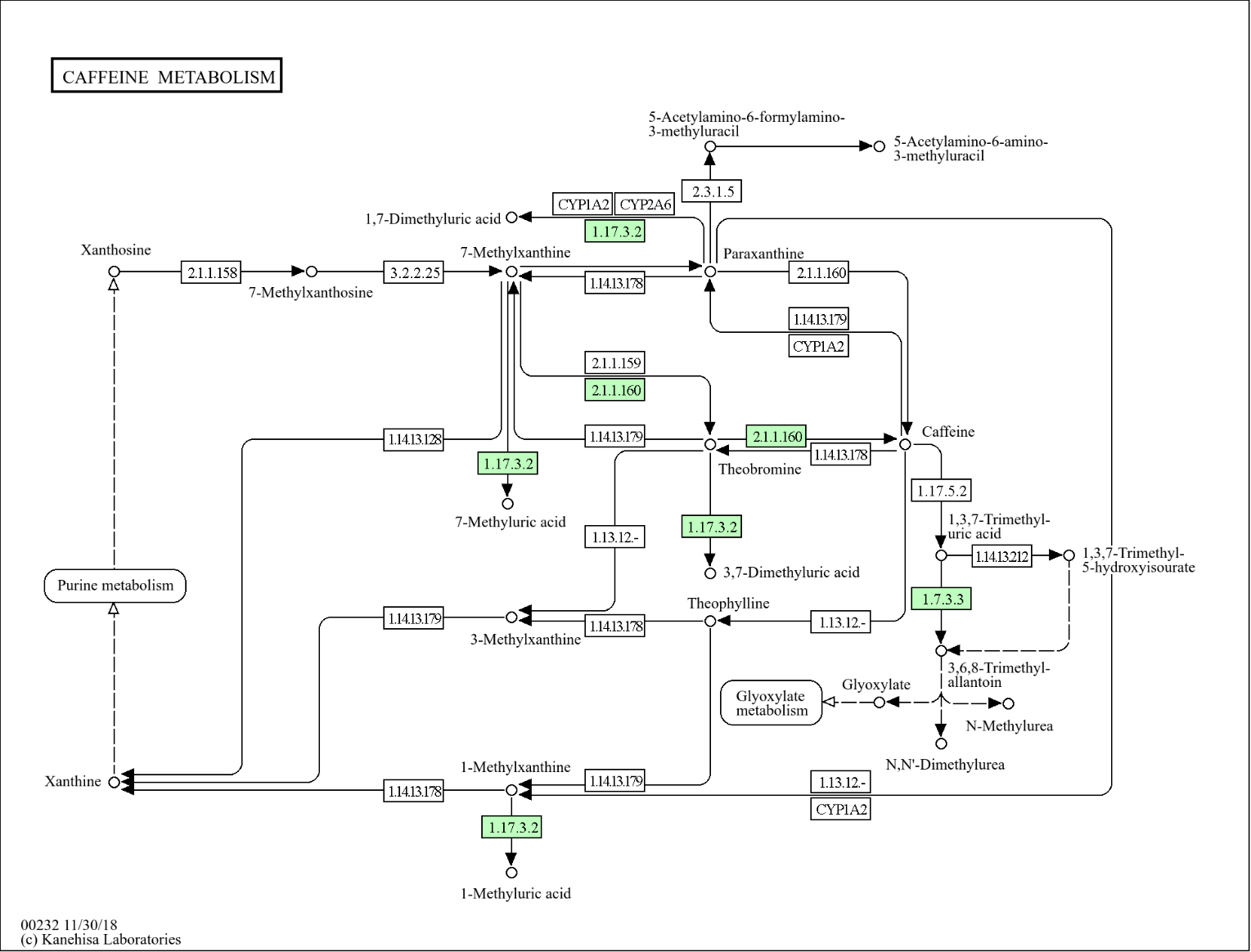
complex methylxanthines.

**Note S3 Figure 7:** KEGG pathway mapper illustration of the reference caffeine metabolism mapping the KofamScan results from the predicted Silene latifolia proteome. Kegg orthologue functions present in the mapped proteome are highlighted green.
